# Modes of action and *in planta* antifungal activity of *Olea europaea* defensin OefDef1.1-derived peptide variant

**DOI:** 10.1101/2025.11.05.686674

**Authors:** Ruby Tiwari, S. Hamsa, James Godwin, Meenakshi Tetorya, Emery Usher, Dilip Shah

## Abstract

Peptide-based biopesticides represent a promising strategy for sustainable disease control in agriculture. Synthetic antifungal peptides incorporating the γ-core motif of plant defensins offer multiple modes of action (MoA) and potential as biofungicides. We investigated a synthetic variant of the olive defensin OefDef1.1 for antifungal activity, structure-function relationships, and MoA against *Botrytis cinerea*, the necrotrophic pathogen causing gray mold. A disulfide-bridged peptide, GMAOe1C_V1*, derived from OefDef1.1 (G32–Y53) and modified with hydrophobic amino acid substitutions, inhibited *B. cinerea* growth *in vitro* and reduced lesion formation in detached leaves. Foliar application of GMAOe1C_V1* suppressed disease symptoms in pepper plants. Mechanistically, GMAOe1C_V1* rapidly permeabilized fungal plasma membranes and accumulated in vacuoles, triggering vacuolar expansion and cell death. It also inhibited protein synthesis *in vitro* and *in vivo*, suggesting a role as a translation inhibitor. Alanine scanning mutagenesis of the non-disulfide bridged variant identified the ^7^RHSKH^11^ motif as essential for antifungal activity. Circular dichroism revealed an unstructured conformation with minimal secondary structure. Transcriptomic analysis of GMAOe1C_V1* treated *B. cinerea* germlings showed downregulation of genes involved in mitochondrial function and amino acid biosynthesis. These findings demonstrate the potential of an olive defensin-derived peptide as a bio-inspired antifungal agent with multifaceted MoA, supporting its development for crop protection.

## INTRODUCTION

Fungi are major agricultural pathogens globally causing billions of dollars in crop yield losses every year (Fones et al., 2020). Plant breeding has long been used to increase fungal resistance in crops and thus improve crop yields. However, many commercially available crop cultivars do not have genetic resistance to economically important fungal pathogens (van Esse et al., 2020). Therefore, growers must rely on costly chemical fungicides to protect crops from fungal diseases (Oliver and Hewitt, 2014). These fungicides have serious repercussions for the environment and even human health. Moreover, evolution of resistance in many fungal pathogens to fungicides with single-site mode of action (MoA) poses a serious and growing threat to crop protection globally (Steinberg and Gurr, 2020). Thus, it would be prudent to look for an alternative strategy for management of fungal diseases. Sustainable agriculture will increasingly depend on the use of eco-friendly non-toxic fungicides. Small synthetic or natural peptides that directly kill fungal pathogens using multiple MoA can be used for a highly effective crop protection system (Leannec-Rialland et al., 2022; Lobo and Boto, 2022; Rosa et al., 2022; Montesinos, 2023).

Plant defensins are a large well-characterized family of cysteine-rich peptides broadly distributed in angiosperms (Cools et al., 2017; Parisi et al., 2019). These peptides are typically 45-54 amino acids in length and contain a conserved pattern of eight cysteine residues. The compact and globular three-dimensional structure of each plant defensin consists of a common fold known as a cysteine-stabilized αβ (CSαβ) fold (Kovaleva et al., 2020). It consists of one α-helix and three antiparallel β strands stabilized by four disulfide bonds which enhance stability against temperature, pH and proteases. Despite their structural conservation, plant defensins show minimal sequence similarity beyond the eight highly conserved cysteine residues permitting discovery of antifungal peptides with varied MoA (Van der Weerden and Anderson, 2013).

Because of their length, plant defensins are not amenable to chemical synthesis to permit extensive characterization of their variants for antifungal activity and MoA. Chemically synthesized truncated defensins containing the functional γ-core motif (GXCX₃-_9_C) overcome this limitation by enabling antifungal screening and *in planta* disease assays. Several defensin-derived peptides have now been characterized for their antimicrobial activity and MoA (Sathoff et al., 2019; Tetorya et al., 2023; Kalunke et al., 2025).

We previously reported a novel gene family encoding highly cationic histidine- and tyrosine-rich defensins in the olive (*Olea europaea*) tree (Li et al., 2019). Members of this gene family were unique to the *Oleaceae* family and were among the highly cationic plant defensins. OefDef1.1, member of this unique defensin family, was previously characterized for its potent antifungal activity *in vitro* against fungal pathogens *Botrytis cinerea* and three *Fusarium* species. This defensin also reduced gray mold disease lesions in the detached leaf assays (Li et al., 2019).

Since the cationic γ-core motif (GACLKNRHSKHYGC) of OefDef1.1 differs in sequence and length from other previously characterized defensins, we obtained a chemically synthesized carboxy-terminal 22-amino acid peptide, designated GMAOe1C_WT containing the γ-core motif but no disulfide bonds and tested it for its antifungal activity *in vitro* against *B. cinerea*. We also designed variants of GMAOe1C_WT with hydrophobic amino acid substitutions and with or without a C3-C21 disulfide bond. GMAOe1C_V1* with a C3-C21 disulfide bond exhibited two-fold higher antifungal activity than GMAOe1C against *B. cinerea*. When applied topically on pepper plants, GMAOe1C_V1* was more effective in controlling gray mold disease *in planta* than the full-length OefDef1.1 and other variants. Further, the RNA-seq analysis identified biochemical and physiological processes affected by the GMAOe1C_V1* challenge in *B. cinerea*.

## RESULTS

### Truncated OefDef1.1-derived peptide variants differ in their antifungal activity

As reported previously, OefDef1.1 is a 53-amino acid defensin rich in histidine and tyrosine. It carries a net charge of +9.75 and 30% hydrophobic residues (Li et al., 2019). It contains a cationic γ-core motif with the sequence GACLKNRHSKHYGC (**Fig. 1A**). We designed a truncated 22-residue variant designated GMAOe1C_WT spanning the γ-core motif carrying a net charge of +4.75 and 27% hydrophobic residues. Based on our previous work with plant defensin MtDef4-derived peptide GMA4CG (Tetorya et al., 2023), we generated two variants GMAOe1C_V1* and GMAOe1C_V2* in which the internal CYC motif was replaced with WFW or FFW, respectively, thus increasing hydrophobic residues to 32%. A C3-C21 disulfide bond was introduced into each variant during chemical synthesis to test the effect of a disulfide bond on the antifungal activity. In addition, GMAOe1C_V1 and GMAOe1C_V2 each lacking a disulfide bond were also synthesized for a comparison of their antifungal activity with their counterparts having a disulfide bond (**Fig. 1A**).

**Fig. 1.**
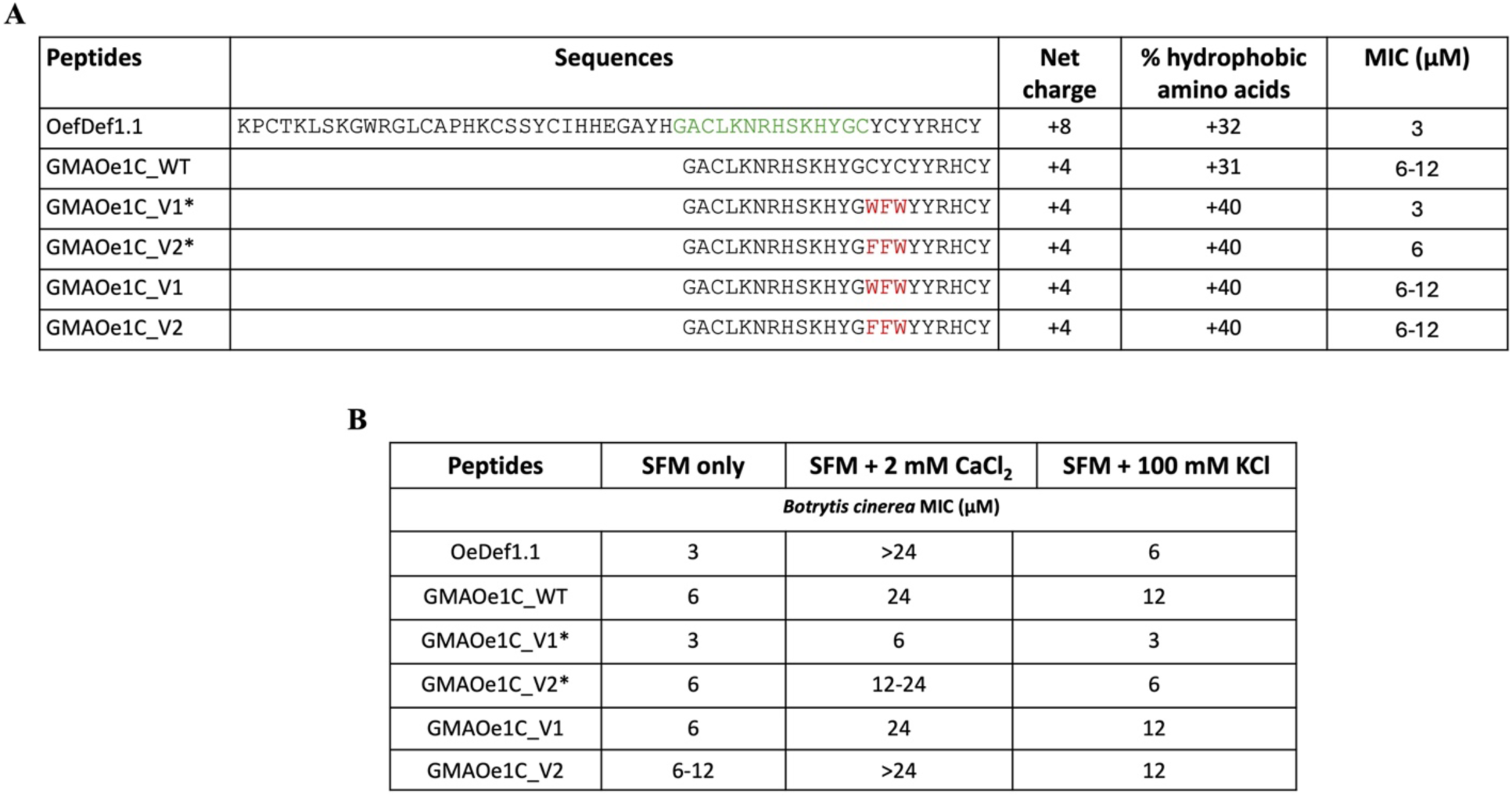
Amino acid sequences, properties and antifungal activity of OefDef1.1-derived GMAOe1C_WT and its variants. **A)** Amino acid sequences of OefDef1.1, OefDef1.1-derived peptide GMAOe1C_WT and its variants. The γ-core motif of OefDef1.1 is shown in green. The amino acid substitutions in the truncated peptide sequences are shown in red. *Asterisks indicate peptide variants with a C3-C21 disulfide bond. The net charge, % hydrophobic amino acids, and minimal inhibitory concentration (MIC) of each peptide against *B. cinerea* are shown. **B)** The MIC of OefDef1.1, GMAOe1C_WT and its variants against *B. cinerea* in SFM supplemented with calcium chloride (CaCl_2_) and potassium chloride (KCl).

Antifungal activity of OefDef1.1, GMAOe1C_V1*, GMAOe1C_V2*, GMAOe1C_V1 and GMAOe1C_V2 against *B. cinerea* was determined in a synthetic fungal medium (SFM). As shown in **Fig. 1A**, all peptides were active but differed in their potency. Whereas OefDef1.1 inhibited this fungus with an MIC value of 3 µM, GMAOe1C_WT containing the C-terminal 22 amino acids of this defensin inhibited this fungus with an MIC value of 6-12 µM. GMAOe1C_V1* and GMAOe1C_V2* had MIC values of 3 and 6 µM, respectively. Thus, GMAOe1C_V1* carrying a WFW substitution and a disulfide bond was as potent as full-length defensin OefDef1.1. The variants lacking the disulfide bond were two- to four-fold less active against *B. cinerea* (**Fig. 1A**). GMAOe1C_V1* peptide was also active against *Colletotrichum gloeosporioides* with the MIC value of 24 µM. It was also previously reported to inhibit *Cercospora sojina* with an MIC value of 3-6 µM (Pokhrel et al., 2025).

Since full-length OefDef1.1 was reported previously to lose its antifungal activity substantially in presence of elevated cations in SFM (Li et al., 2019), *in vitro* antifungal activity of the truncated peptides against *B. cinerea* was tested in SFM supplemented with the physiologically relevant concentrations of 2 mM CaCl_2_ and 100 mM KCl in the plant tissues. In presence of 2 mM CaCl_2_ added to SFM, GMAOe1C_V1* lost its antifungal activity only two-fold with an MIC of 6 µM, whereas all other peptides had MIC values of greater than 12 µM. However, in presence of 100 mM KCl added to SFM, all peptides except GMAOe1C_V1* and GMAOe1C_V2* lost their antifungal activity two-fold (**Fig. 1B**). Thus, GMAOe1C_V1* was the most potent peptide against *B. cinerea* in our *in vitro* antifungal assays.

### Structure prediction using circular dichroism (CD) and AlphaFold2 (AF2)

To understand the structural features of OefDef1.1 and its peptide derivatives, we used CD and structure prediction using AF2. The CD spectrum of the full-length OefDef1.1 revealed structural features consistent with that of a primarily folded protein with α-helical and β-sheet character (Supplementary **Fig. S1A**, solid line). Deconvolution of the secondary structure contributions to the CD spectrum (see Materials and Methods) yielded an estimated 47% β-strand, 4% α-helix, and 49% turn, which seems to underestimate the helix content as predicted by AF2. For direct comparison between spectra, the CD spectrum calculated from the AF2 model is shown in Supplementary **Fig. S1A** as a dashed line. Indeed, the calculated spectrum shows a distinct minimum at 222 nm, which suggests a higher fraction of helix than that of the measured protein spectrum. A possible interpretation for the low helical signal in our measurements is that peptide dynamics may allow the helix to fluctuate in solution.

We previously published the high-quality AF2 generated structure of the full-length OefDef1.1 defensin showing the presence of a cysteine-stabilized α/β motif comprising one α-helix stabilized through tetradisulfide array (Cys3-Cys52, Cys14-Cys34, Cys19-Cys45, and Cys23-Cys47) to three anti-parallel β-strands (Li et al., 2019) (Supplementary **Fig. S1B**). The electrostatic surface representation of this structure revealed cationic amino acids on the surface of the OefDef1.1 structure that are presumably involved in binding to the negatively charged fungal membranes (Supplementary **Fig. S1C**).

The CD analysis showed GMAOe1C_WT, GMAOe1C_V1*, GMAOe1C_V2* peptides to be primarily unstructured in solution (Supplementary **Fig. S2A-C**). The signal in the spectrum of each peptide decreased toward 200 nm which is characteristic of an unstructured peptide. Each peptide has a positive peak around 230 nm which likely arises from the electronic interactions of the aromatic side chains in the sequence; hence, the 230 nm peak is more pronounced in the peptide variants that add aromatic residues, especially Trp (Grishina and Woody, 1994). The AF2 predictions for all three peptides are fairly low-confidence with some short elements of structure consistent with a largely unstructured conformation, likely stabilized by disulfide bridges formed by the remaining cysteine residues (Supplementary **Fig. S2A-C**).

### Identification of the sequence motif responsible for antifungal activity of GMAOe1C_V1

To identify the amino acid sequence motif that governs the antifungal activity of GMAOe1C_V1, alanine scanning mutagenesis of the peptide was performed. Four Ala scanning variants, GMAOe1C_V1_Ala1, GMAOe1C_V1_Ala2, GMAOe1C_V1_Ala3, GMAOe1C_V1_Ala4, were chemically synthesized each carrying substitutions of five amino acids while retaining the two Cys residues (**Fig. 2A)**. The antifungal activity of these variants against *B. cinerea* was compared with the parental GMAOe1C_V1 peptide (**Fig. 2B**). GMAOe1C_V1_Ala1, GMAOe1C_V1_Ala3, GMAOe1C_V1_Ala4 variants showed two-fold reduction in antifungal activity; however, the antifungal activity of GMAOe1C_V1_Ala2 variant showed no activity even at 24 μM. This infers that the amino acid sequence ^7^RHSKH^11^ within the γ-core motif is crucial for the antifungal activity of GMAOe1C_V1 (**Fig. 2C)**.

**Fig. 2.**
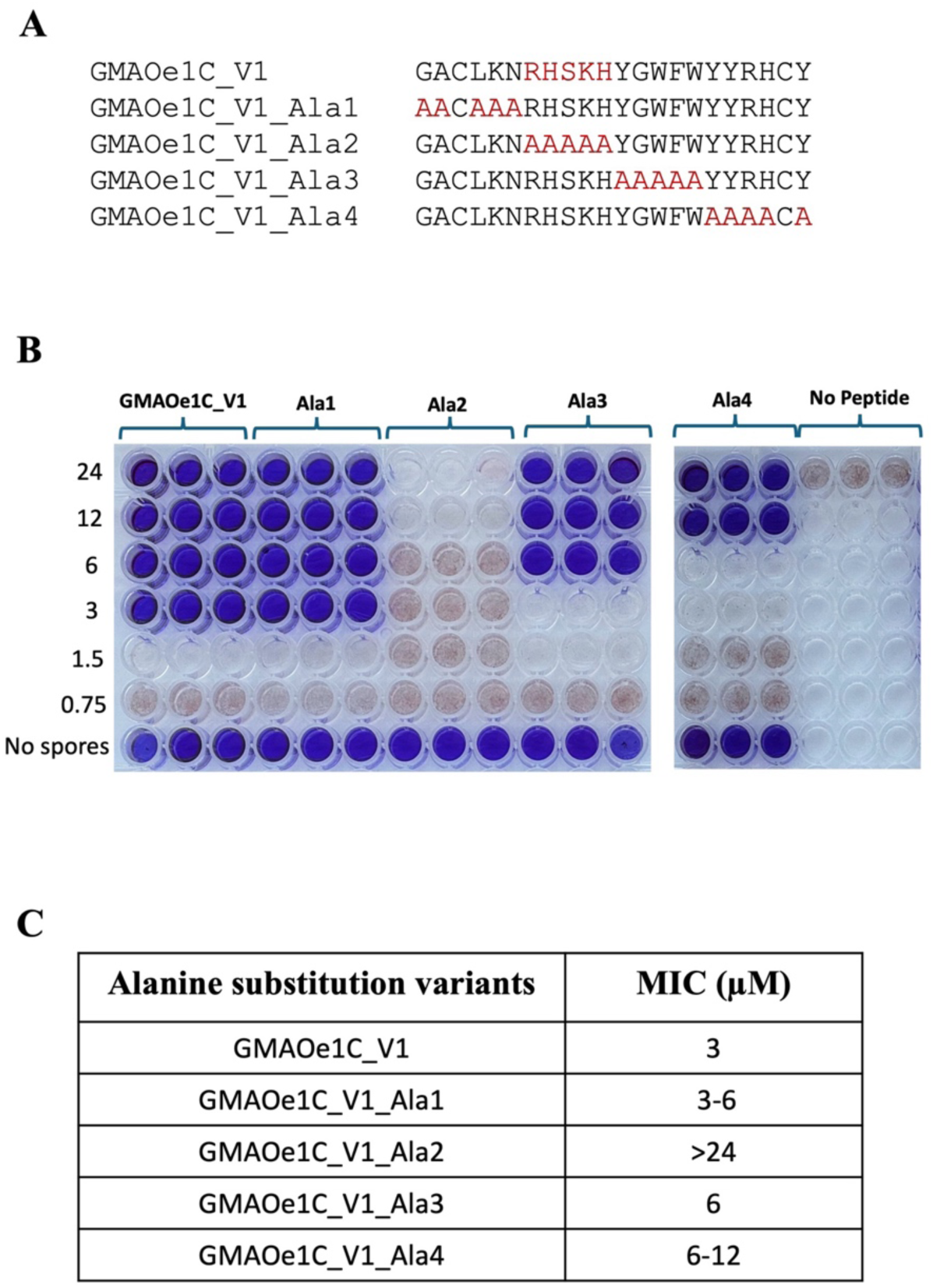
Identification of the active motif required for antifungal activity of GMAOe1C_V1 against *B. cinerea*. **A)** Amino acid sequences of the alanine (Ala) scanning variants of GMAOe1C_V1. **B)** Resazurin cell viability assay to determine the MIC value of each alanine scanning variant. **C)** MIC values of variants determined via resazurin assay. Values were obtained from three independent biological replicates.

### OefDef1.1 and OeDef1.1-derived peptides reduce gray mold symptoms in detached pepper leaves

Since OefDef1.1, GMAOe1C_WT, GMAOe1C_V1*, GMAOe1C_V2*, GMAOe1C_V1 and GMAOe1C_V2 were all active against *B. cinerea in vitro*, we investigated their ability to reduce gray mold symptoms when applied on detached pepper leaves. Three different concentrations (1.5, 3, and 6 μM) of all five peptides were applied on the surface of pepper leaves followed by *B. cinerea* conidia as shown in Supplementary **Fig. S3A** and the gray mold disease lesions were observed three days post-inoculation (dpi) (Supplementary **Fig. S3B-3G**). The CropReporter images showed large lesions on leaves treated with no peptide. All peptides failed to reduce disease lesions when used at a sublethal concentration of 1.5 μM (Supplementary **Fig. S3B, E**). However, only OefDef1.1 and GMAOe1C_V1* significantly reduced lesion sizes at 3 μM (Supplementary **Fig. S3C, F)**. Similarly, GMAOe1C_V2* was effective at 6 μM, which correlated with its antifungal activity *in vitro* (Supplementary **Fig. S3D, G**). The quantification of lesion sizes using ImageJ software confirmed that GMAOe1C_V1* was most effective among the OefDef1.1-derived peptides in reducing gray mold symptoms on pepper leaves (Supplementary **Fig. S3C-D, F-G**).

### OefDef1.1 and OefDef1.1-derived peptides reduce gray mold disease symptoms *in planta*

The *in vitro* and detached leaf antifungal assays revealed that OefDef1.1 and GMAOe1C_V1* peptides were highly effective against *B. cinerea*. Therefore, we tested the potential of these peptides to control gray mold disease in pepper plants. We also tested GMAOe1C_WT and GMAOe1C_V2* peptides in this study for comparison (**Fig. 3**). Four-week-old pepper plants spray-inoculated with *B. cinerea* followed by peptide spray at the concentrations of 6 and 12 μM showed better tolerance to gray mold disease as compared to the no peptide control. Among all peptides tested, the CropReporter images showed that GMAOe1C_V1* applied at 12 μM has the best post-inoculation ability to control gray mold *in planta* (**Fig. 3A**). In addition, the visual symptoms were scored manually (Segarra et al., 2007) and plotted, which indicated that GMAOe1C_V1* was most efficient in conferring post-inoculation antifungal activity against *B. cinerea* in pepper plants (**Fig. 3B**).

**Fig. 3.**
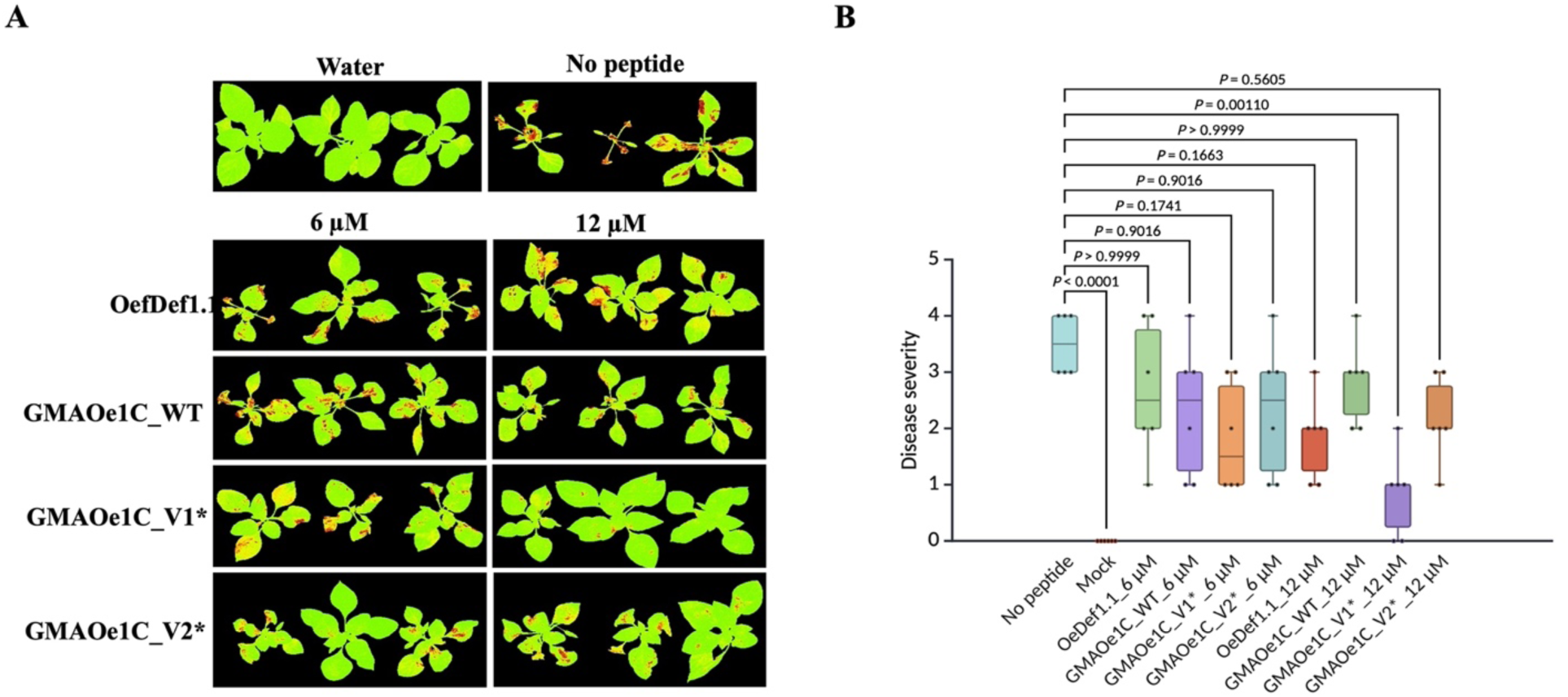
Post-inoculation antifungal activity of OefDef1.1 and OefDef1.1-derived peptides against the gray mold disease in pepper plants. **A)** For post-inoculation antifungal activity assay, four-week-old pepper plants were sprayed with 1 mL of 5 x 10^5^ spores/mL followed 2 h later by 2 mL of each peptide OefDef1.1, GMAOe1C_WT, GMAOe1C_V1*, GMAOe1C_V2* at a concentration of 6 and 12 μM. The CropReporter images were taken at 3 dpi. **B)** Quantification of gray mold symptoms was done as described by (Segarra et al., 2007). Statistical analysis was done using Welch’s one-way ANOVA with Games-Howell multiple comparisons test used. ns-p>0.05, p<0.05, p<0.01, p<0.001, p<0.0001.

### GMAOe1C_V1* permeabilizes plasma membrane of *B. cinerea* germlings and is internalized into germling cells

To determine if GMAOe1C_V1* was capable of permeabilizing the plasma membrane of *B. cinerea* germlings, time-lapse confocal microscopy was performed to study the uptake of SYTOX Green (SG) dye at a sublethal dose of 1.5 μM GMAOe1C_V1*. The SG dye was seen to gain entry from the tip of *B. cinerea* germlings and reach the conidial head. SG signal was observed in germlings within 1 min, and stronger signal was observed within 10 min in the nuclei of the apical germ tube cell (**Fig. 4A, Video 1**). Thus, GMAOe1C_V1* disrupts the plasma membrane of *B. cinerea* germlings within minutes.

**Fig. 4.**
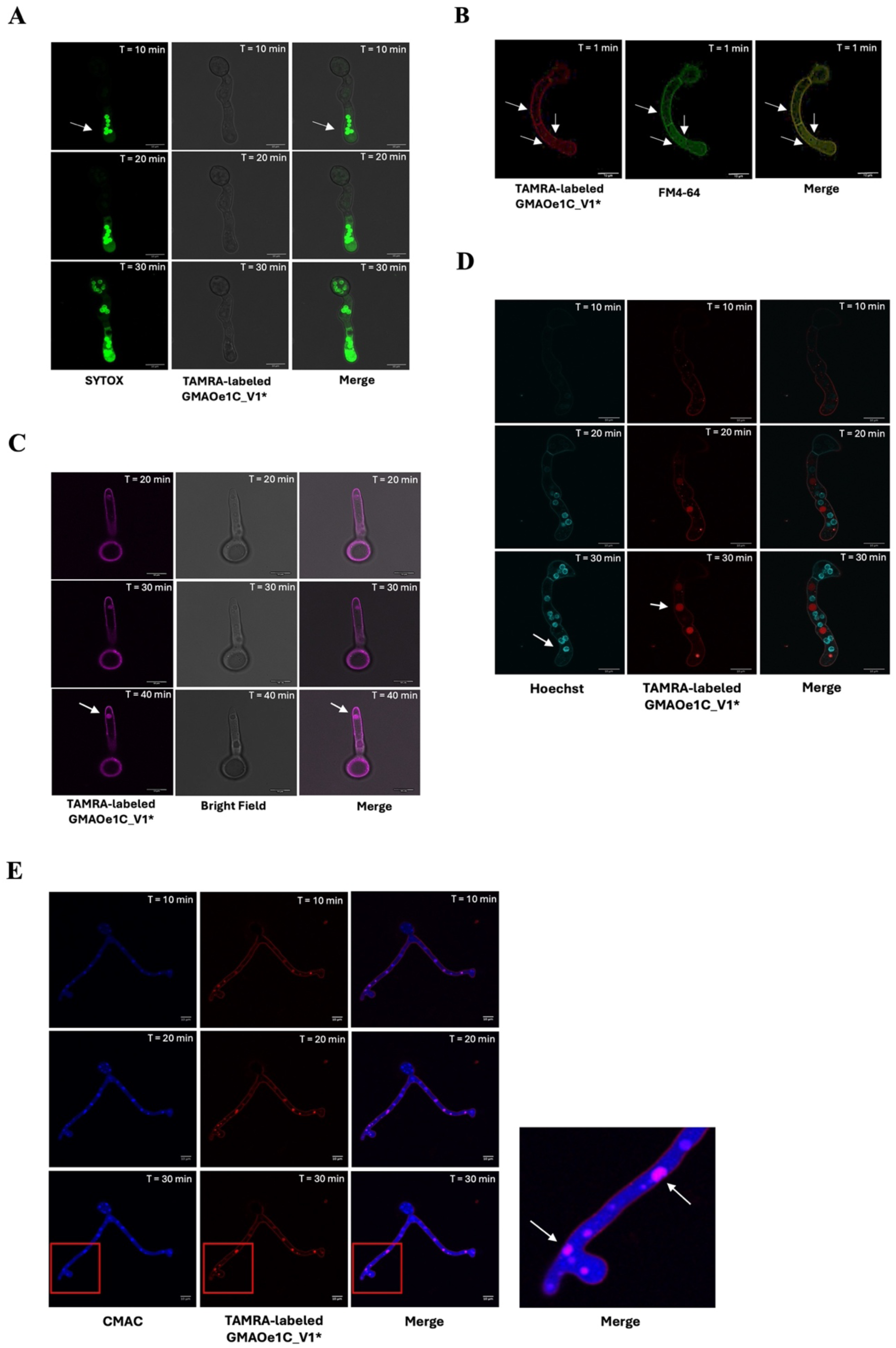
GMAOe1c_V1* induces membrane permeabilization and is internalized in *B. cinerea* germlings. **A)** Time-lapse images of *B. cinerea* germlings incubated with SYTOX Green (SG) dye and GMAOe1C_V1* (1.5 μM). The images show an increase in the internalization of SG dye with time (arrow indicates entry of SG into the germlings from the tip). B) Tetramethyl rhodamine (TAMRA)-labeled GMAOe1C_V1* is internalized in *B. cinerea* germlings through multiple foci (arrow indicates entry spots of TAMRA-labeled GMAOe1C_V1* (red signal), FM4-6 (green signal) and merge of both signals (yellow signal). **C**) Time-lapse images of *B. cinerea* germlings incubated with TAMRA-labeled GMAOe1C_V1* (arrow indicates accumulation of TAMRA-labeled GMAOe1C_V1* in vacuoles). D) Nuclear DNA-specific Hoechst33258 determined that TAMRA-GMAOe1C_V1* (3 μM) does not colocalize with the nucleus (blue signal indicated with arrow) but accumulates in vacuoles (red signal indicated with arrow). The size of vacuoles increased in a time-dependent manner. **E)** Confocal images showing the subcellular localization of TAMRA-GMAOe1C_V1* (6 μM) in vacuoles using the vacuole-specific CMAC dye. Arrow indicates merge of CMAC and TAMRA-labeled GMAOe1C_V1* signals as pink). Scale bar = 10 μm.

Next, we used tetramethyl rhodamine (TAMRA) labeled-GMAOe1C_V1* to determine its ability to gain entry into germling cells of *B. cinerea* and identify its subcellular location within the cells using the time-lapse confocal microscopy. The MIC of TAMRA-labeled GMAOe1C_V1* was 6 μM, two-fold higher than that of unlabeled GMAOe1C_V1*. To explore the entry spots of TAMRA-labeled GMAOe1C_V1*, germlings were treated with the labeled-peptide at a concentration of 1.5 μM in the presence of cell membrane and endocytosis-specific dye, FM4-64 (10 μM). The labeled peptide co-localized with FM4-64 dye indicating binding to the plasma membrane at 1 min post-challenge (**Fig. 4B**). GMAOe1C_V1* gradually accumulated at putative entry spots at the cell surface of *B. cinerea* germlings (shown in yellow at 1 min) and entered the germlings via multiple foci (**Fig. 4B, Video 2**).

### Subcellular localization of GMAOe1C_V1* to the vacuoles in *B. cinerea* germlings

The internalization of the TAMRA-labeled GMAOe1C_V1* in germlings was also tracked at a lethal dose of 6 µM which showed that the labeled peptide gradually accumulated in a specific hotspot in cytoplasm at 30-40 min post-challenge (**Fig. 4C, Video 3**). To determine if TAMRA-labeled GMAOe1C_V1* was targeted to specific organelles, a vacuole-specific dye CMAC and nuclear staining dye Hoechst 33258 were used. Notably, TAMRA-labeled peptide (3 μM) did not show any colocalization with the nucleus (**Fig. 4D, Video 4**). However, it was noted that TAMRA-labeled GMAOe1C_V1* (6 μM) gained entry and started accumulating in the vacuoles. The gradual accumulation of the peptide in the vacuoles led to vacuole expansion and the collapse of germlings (**Fig. 4E, Video 5)**. Moreover, propidium iodide (PI) staining and bright field images indicated that germlings started to collapse in 15 min upon challenge with unlabeled peptide, i.e. GMAOe1C_V1 (3 μM). The PI stain spread and stained the whole germlings 30 min post-challenge, indicating cell death (Supplementary **Fig. S4**).

### GMAOe1C_V1* inhibits protein synthesis *in vitro* and *in vivo* but does not bind to ribosomal RNA

Some antifungal peptides inhibit protein translation *in vitro* (Godwin et al., 2024; Li et al., 2024; Kalunke et al., 2025) and *in vivo* (Godwin and Shah, 2025). To test protein translation inhibitory activity of GMAOe1c_V1*, an eukaryotic wheat germ extract *in vitro* translation inhibition system was utilized. The luciferase reporter transcripts were produced *in vitro* and translated to luciferase in the presence of different concentrations of GMAOe1C_V1*. The study showed that the translation inhibitory activity of GMAOe1C_V1* was concentration-dependent (**Fig. 5A**). When compared with enzyme activity in presence of water, luciferase activity was reduced by only 50% at a higher peptide concentration of 24 µM and 70% at a concentration of 48 µM. Cycloheximide, an eukaryotic translation inhibitor, was used as a positive control where no luciferase activity was detected.

**Fig. 5.**
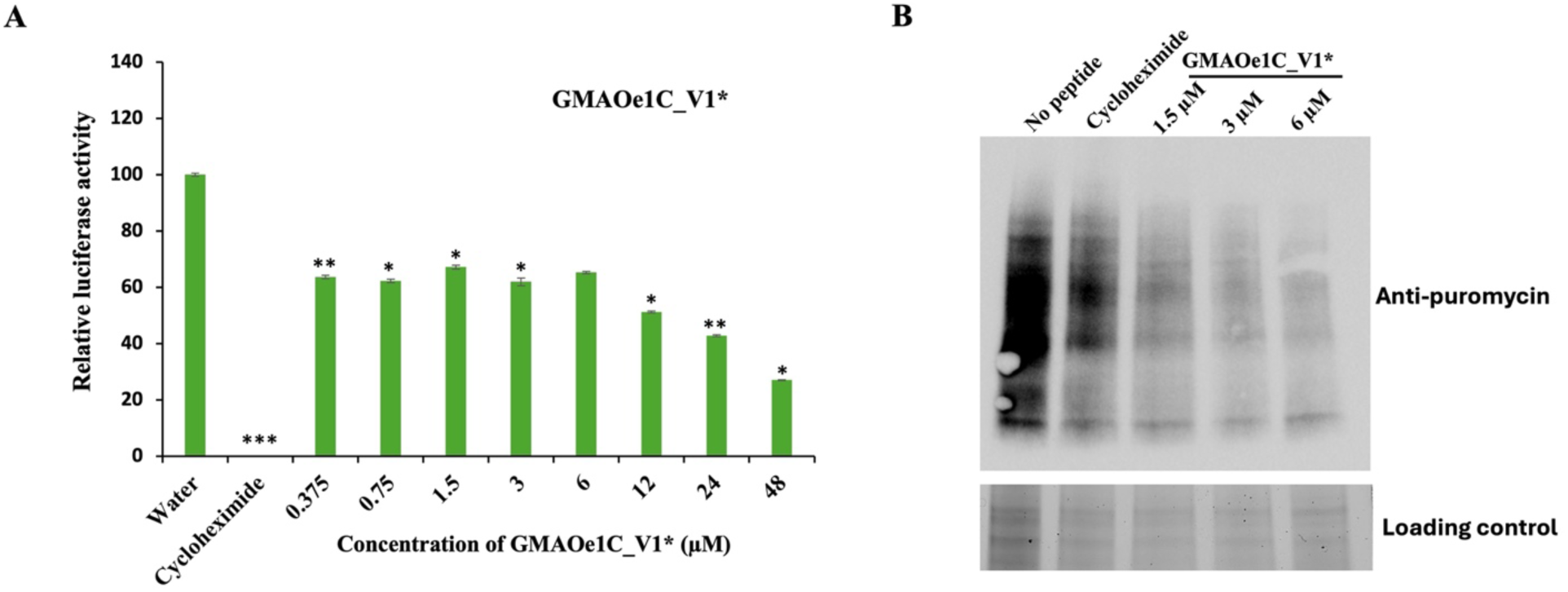
*In vitro* and *in vivo* translation inhibition activity of GMAOe1C_V1*. **A)** The luciferase activity of GMAOe1C_V1* was quantified. Sterile water was used as a negative control, and the translation activity was considered as 100%. Cycloheximide (100 μg/mL) was used as positive control. Student’s t-test was used to perform statistical analysis, (Biological replicates, *n* = 2, \*\**p* < 0.01, \*\*\**p* < 0.001, ns, *p* > 0.05). Bars = mean ± SE. **B)** *In vivo* puromycin labeling showed translation inhibition activity of GMAOe1C_V1* (1.5, 3 and 6 μM).

Next, we determined if GMAOe1C_V1* was capable of inhibiting protein translation *in vivo* in fungal cells using puromycin labeling (Godwin and Shah, 2025). *B. cinerea* germlings were treated with GMAOe1C_V1* at three different concentrations of 1.5, 3 and 6 μM to assess the inhibition of protein translation *in vivo*. The study revealed that the protein synthesis in *B. cinerea* germlings was reduced on treatment with lethal or sub-lethal doses of GMAOe1C_V1* (**Fig. 5B**). One way the peptide could inhibit protein synthesis is by binding to ribosomal RNA or proteins (Li et al., 2024). Therefore, fungal RNA was incubated with GMAOe1C_V1* at concentrations ranging from 3 to 48 μM. NCR13_PFV1 peptide was used as a positive control (Godwin et al., 2024). Whereas NCR13_PFV1 peptide bound to fungal rRNA, GMAOe1C_V1* peptide did not show any binding, indicating different MoA of GMAOe1C_V1* for inhibition of protein synthesis (**Fig. S5**).

### GMAOe1C_V1* treatment induces transcriptomic reprogramming in *B. cinerea*

To elucidate molecular mechanisms underlying antifungal activity of GMAOe1C_V1* against *B. cinerea*, RNA-Seq was performed to analyze global transcriptional alterations following peptide treatment. Considering the *in vitro* inhibitory and cell death inducing effects of GMAOe1C_V1* in *B. cinerea* germlings, a lethal dose of 3 μM was chosen and transcriptome analysis was conducted at two-time points; 30 min and 1 h following peptide treatment with respect to non-treated 0 h control (Supplementary **Fig. S6A).** Principal Component Analysis (PCA) plot revealed separate clustering of peptide treated germlings (30 min, 1h) from the control group (0 h) along PC1 (40%), indicating differences in gene expression profiles in *B. cinerea* in response to GMAOe1C_V1* challenge (Supplementary **Fig. S6B).**

Upon GMAOe1C_V1* treatment, a combined total of 967 genes were differentially expressed as compared to the control group. Among the 967 DEGs, 310 and 506 genes were upregulated while 36 and 115 genes were downregulated at 30 min and 1 h, respectively **(Fig. 6A, B**; Supplementary **Fig. S6C,** Supplementary **Fig. S7)**. The peptide treatment led to a time-dependent increase of differentially expressed genes in *B. cinerea*. A total of 238 DEGs (219 upregulated and 19 downregulated) were identified as common to both time points **(Fig. 6A)**. To validate the results of RNA-Seq, a subset of DEGs from both time points was selected. In response to GMAOe1C_V1* treatment, Bcin05g00040, Bcin01g03760, Bcin05g00740, Bcin03g00690, Bcin01g03350, Bcin12g02040 showed upregulation at either 30 min or both time points (30 min and 1h). Two genes, Bcin14g04580 and Bcin11g04710 showed downregulation at both timepoints as observed in our RNA-Seq data. Consistent with the RNA-seq data, a subset of 3 genes, Bcin13g02010, Bcin09g03670, and Bcin01g05680 exhibited upregulation at only 1h post peptide treatment (Supplementary **Fig. S8;** Supplementary **Table S3**).

**Fig. 6.**
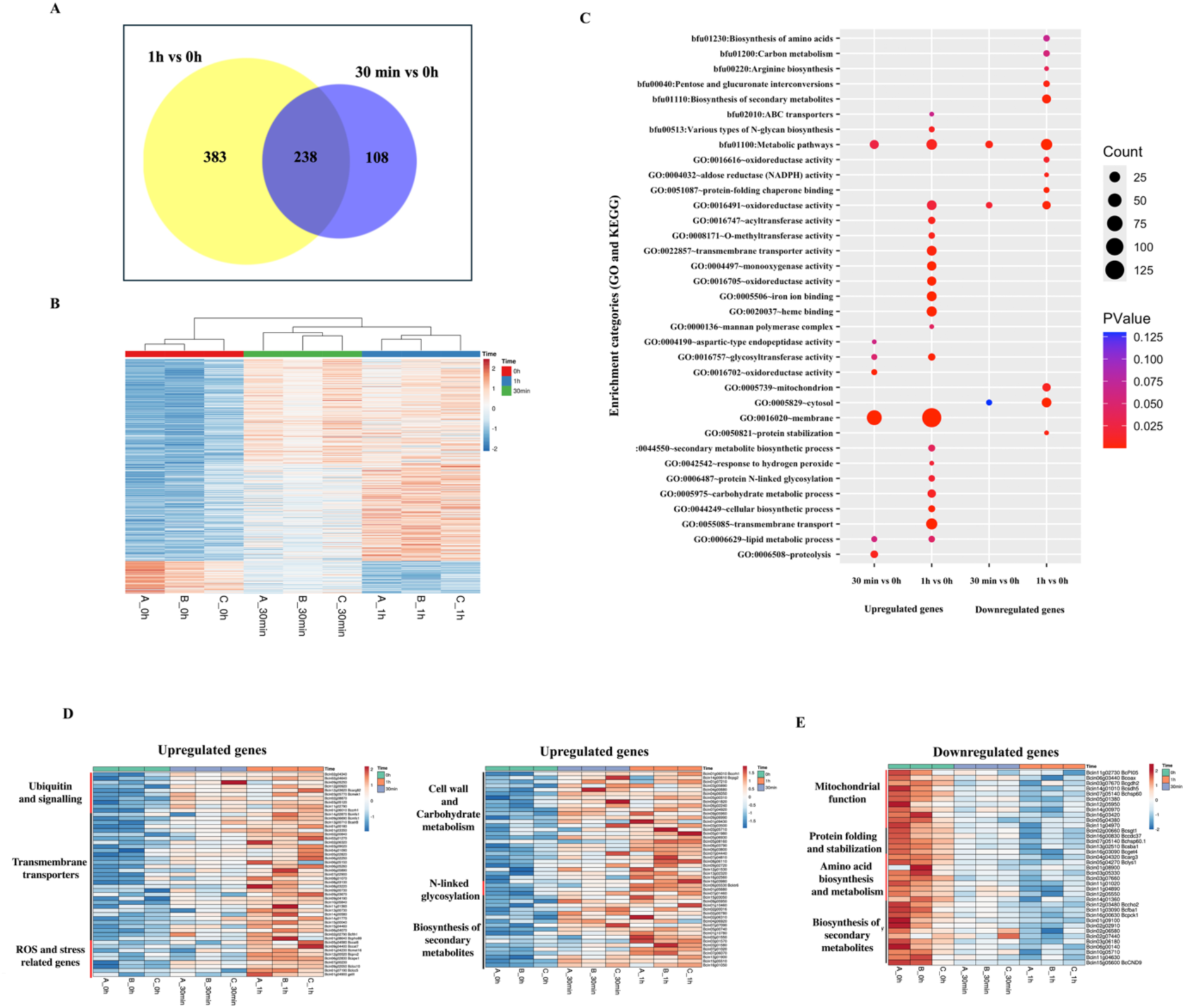
Transcriptomic reprogramming in *B. cinerea* upon GMAOe1C_V1* treatment. **A)** Venn diagram showing the number of exclusive and shared Differentially Expressed Genes (DEGs) at 30 min and 1 h post GMAOe1C_V1* treatment. Genes were classified as differentially regulated if they exhibited a log2 fold change of ≥1 for upregulated genes and ≤ -1 for downregulated genes in comparison to the control (0 h), with a p-adjusted value of ≤ 0.05 following the False Discovery Rate (FDR) correction. **B)** Heatmap depicting hierarchical clustering and expression patterns of all DEGs for each sample based on the variance stabilized counts of all genes from three biological replicates. **C)** Gene Ontology and KEGG enrichment analysis for functional characterization of DEGs upon GMAOe1C_V1* treatment. The dot plots depict the number of genes (circle size) and significance by p-value (colors) for top enriched functional GO and KEGG categories in both upregulated and downregulated DEGs. **D, E)** Heatmaps representing the relative expression of Upregulated **(D)** and Downregulated **(E)** genes related to various functional categories. Each horizontal row represents one gene, and vertical columns represent samples from all three biological replicates. The variance stabilized counts generated from DESeq2 were used for the generation of all heatmaps.

To understand the target processes and pathways influenced by GMAOe1C_V1* challenge, GO and KEGG enrichment analysis were performed. The upregulated DEGs were related to transmembrane transporters, carbohydrate and lipid metabolic processes, response to reactive oxygen species (ROS) and proteolysis. Interestingly, the downregulated DEGs were exclusively enriched for genes involved in mitochondrial function under the cellular component category and pathways like biosynthesis of amino acids and pentose and glucuronate interconversions (**Fig. 6C**; Supplementary **Fig. S9-S10;** Supplementary **Table S4).**

### GMAOe1C_V1* treatment alters expression of key stress and metabolic genes in *B. cinerea*

Antifungal peptides have been reported to trigger fungal signaling cascades that induce ROS generation, membrane permeabilization, and cellular disorganization causing peptide-induced oxidative damage and membrane disruption as key components of its antifungal mechanism (Vriens et al., 2014). In our data, multiple genes associated with proteolysis, autophagy and ubiquitination - specifically aspartic (*Bcap1*, *Bcap4*, *Bcap8*) and serine endopeptidases, *Bcser1* (Bcin10g02530), Bcin02g0460 (ubiquitin-3-binding domain) exhibited significant upregulation **(Fig. 6D)** (Supplementary **Table S5**). GMAOe1C_V1* treatment activated genes associated with the general stress response, specifically those involved in signal transduction and antioxidant mechanisms **(Fig. 6D)**. In response to oxidative stress induced by peptide treatment, genes encoding glutathione S-transferase (*Bcmet16*, Bcin07g00230, *gstII*), catalase (*Bccat6*, *Bccat7*), and cupredoxin (*Bclcc10*, *Bclcc5*) were activated significantly (Supplementary **Table S5**). In *B. cinerea*, membrane transporters play key roles in mediating multi-drug resistance to a wide range of antifungal agents by actively extruding toxic compounds, thereby increasing pathogen tolerance and survival (Hayashi et al., 2002; Stefanato et al., 2009; Sofianos et al., 2024). In our analysis, over 30 transmembrane transporter-related genes including MFS transporter *Bcmfs1*, ABC transporter *BcatrB*, iron permease *Bcfth1*, Bcin01g03550, were activated (**Fig. 6D**). The genes related to cell wall organization in fungi, specifically those involved in cellulose and chitin synthesis, cell wall degrading enzymes, and N-protein glycosylation, were found to be upregulated. The N-linked protein glycosylation process is crucial for maintaining cell wall integrity in response to antifungal peptides (Plaza et al., 2025). Genes associated with this process: *Bcktr6* (Bcin06g05530), Bcin09g05950, and Bcin15g03050 encoding alpha-1,6-mannosyltransferase were activated in response to peptide treatment (**Supplementary Table S5**). Together, the upregulation of genes involved in protein degradation, redox stress, membrane remodeling and cell wall integrity suggests that the peptide exerts multifaceted stress on fungal cells.

This study further identified downregulated transcripts associated with amino acid biosynthesis and gluconeogenesis, which are precursors for interconnected metabolic pathways relevant to energy metabolism and mitochondrial respiration in fungi **(Fig. 6E**; Supplementary **Fig. S9**). The DEGs related to amino acid synthesis and metabolism, specifically within the glycine, serine, arginine, tyrosine and lysine pathways, were impacted **(Fig. 6E**; Supplementary **Fig. S11;** Supplementary **Table S6)**. DEGs including the arginine biosynthesis gene *Bcarg3* (Bcin04g04320), the lysine biosynthesis gene *Bclys1* (Bcin05g04270) and a gene associated with kynureninase activity in tryptophan metabolism (Bcin03g07660) exhibited downregulation post-peptide treatment **(Fig. 6E)**. Genes associated with mitochondrial processes, specifically those involved in sulfur metabolism, the electron transport chain, alternative respiration, and ergosterol biosynthesis, such as Bcin12g05950, *Bcgdh2* (Bcin03g07670), Bcin16g03420, *Bcsdh5* (Bcin14g01010), *Bcoax* (Bcin06g03440), *BcPIO5* (Bcin11g02730) and Bcin05g01380, exhibited significant downregulation **(Fig 6E**; Supplementary **Table S6)**. Additionally, transcripts associated with gluconeogenesis, specifically *Bcfba1* (Bcin11g03090) and *Bcpck1* (Bcin16g00630), and Bcin14g04580, critical for virulence and pathogenesis, were also affected by peptide treatment (Liu et al., 2018). (Supplementary **Table S6**). The peptide’s effect on these pathways suggests that it not only inhibits the growth of *B. cinerea* but may also impair the pathogen’s metabolism and virulence mechanisms, thus enhancing its antifungal efficacy. The identified transcriptional changes following peptide treatment likely indicate compensating mechanisms or damage control responses, implying that the antifungal efficacy of GMAOe1C_V1* may arise from its ability to simultaneously interfere with several critical cellular processes.

## DISCUSSION

Plant defensins are a large, well characterized family of antimicrobial peptides that have been extensively probed for their antifungal activity against fungal pathogens. Although the presence of four disulfide bonds makes them highly stable to proteolysis, high temperature and pH, plant defensins are not amenable to chemical synthesis. Therefore, screening of variants for increased antifungal activity and varied MoA has been difficult. Synthetic defensin-derived γ-core motif peptides with potent antifungal activity have caught attention as promising starting templates for designing newer sustainable peptide-based fungicides. OefDef1.1 is a unique His- and Tyr-rich cationic defensin from the perennial woody olive tree characterized for its broad-spectrum antifungal activity *in vitro* and semi-*in planta* and multiple MoA (Li et al., 2019). It contains a γ-core motif with the sequence GACLKNRHSKHYGC, different from previously reported γ-core motif sequences of plant defensins. Here, we used an OefDef1.1-derived carboxy-terminal 22-residue peptide GMAOe1C_WT containing the 14-residue γ-core motif to probe its *in vitro* antifungal activity against *B. cinerea*, a destructive necrotrophic pathogen. In addition, we generated variants of this peptide and identified a variant which displayed superior antifungal activity, further elucidated MoA and probed its potential as a spray-on fungicide for control of the gray mold disease in pepper. Our findings provide an additional example of a peptide-based bio-fungicide distinct from the canonical and conventional fungicides in use today. However, it should be noted that it may be necessary to further optimize its sequence and stability, produce this peptide at scale cost-effectively and explore its synergy in combination with the conventional fungicides (Rosa et al., 2022).

The truncated peptides, GMAOe1C_WT inhibited the growth of *B. cinerea in vitro* with an MIC of 6-12 µM. We designed two additional variants of this peptide, designated GMAOe1C_V1* and GMAOe1C_V2*, in which the CYC motif was replaced with WFW and FFW, respectively. Further, a chemically synthesized disulfide bond (C3-C21) was introduced into each of these peptides making each peptide pseudo-cyclic and likely more stable. For assessing the importance of a disulfide bond for antifungal activity, two other variants, GMAOe1C_V1 and GMAOe1C_V2, were synthesized each lacking a disulfide bond. Antifungal activity of these peptides revealed that GMAOe1C_V1* containing the replacement of CYC with WFW motif and a disulfide bond exhibited 2 to 4-fold more potent antifungal activity than GMAOe1C_V2* which contained FFW motif. Thus, the presence of two Trp residues contributes to enhanced antifungal activity as in plant defensin MtDef4- and MtDef5-derived peptides (Tetorya et al., 2023; Kalunke et al., 2025). Further, addition of a disulfide bond also contributed to the enhanced antifungal activity of GMAOe1C_V1* against *B. cinerea*. It is likely that a pseudo-cyclic GMAOe1C_V1* is more stable inside the cells of this pathogen than its linear counterpart. It is also worth noting that GMAOe1C_V1* retains its potent antifungal activity in the presence of elevated cations in the fungal growth medium, whereas other variants lose their antifungal activity substantially further indicating the potential of GMAOe1C_V1* as a promising antifungal peptide. The biochemical basis for the cation-tolerant antifungal activity of this peptide remains to be determined. It will be interesting to determine if this peptide still binds to the cell wall of conidia and germlings in the fungal growth medium with elevated cations. Further, it will be informative to determine interaction of this peptide with the fungal cell wall polysaccharides such as β-glucans and chitin (Bleackley et al., 2019). GMAOe1C_V1* when spray-applied on pepper plants was also more effective in conferring tolerance to *B. cinerea* than other peptides used in this study. Greater cation-tolerant antifungal activity *in vitro* of this peptide than that of other GMAOe1C peptides might be a contributing factor to its greater *in planta* antifungal efficacy (Li et al., 2019; Li et al., 2024).

The structural analysis of GMAOe1C_WT and its variants using CD and AlphaFold2 analyses revealed that they were all primarily unstructured in solution with some short element of structure in sequences carrying Trp and Phe amino acid substitutions. Overall, the CD spectra indicated the dynamic nature of short peptides. The dynamic nature of these peptides may have several benefits over structured peptides in affording greater flexibility to adopt multiple conformations and ability to interact with a variety of intercellular targets. The disordered structure can contribute to its broad-spectrum antimicrobial activity. The Alanine scanning mutagenesis of the GMAOe1C_V1 peptide identified ^9^RHSKH^13^ as the most critical sequence motif important for antifungal activity against *B. cinerea*. This sequence is mostly composed of positively charged amino acids. In addition, GMAOe1C_Ala_V3 and GMAOe1C_Ala_V4 had 2- and 4-fold reduction in antifungal activity, respectively, indicating the importance of hydrophobic aromatic amino acids. These alanine scanning mutants of GMAOe1C_V1 will be useful in future studies to further elucidate the structure-activity relationships and MoA of this peptide.

Unlike chemical fungicides that target a specific enzyme or protein, GMAOe1C_V1* does not have a single biochemical target in the germling cells of *B. cinerea*. The MoA of this peptide appear to be similar to those of our previously reported plant defensin MtDef4-derived peptide variant GMA4CG_V6 (Tetorya et al., 2023). Like GMA4CG_V6, GMAOe1C_V1* permeabilized the plasma membrane of germling cells within minutes. The time-lapse imaging of *B. cinerea* germlings treated with the fluorescently labeled peptide showed that the peptide was localized on the periphery of the germling, which allowed entry of the peptide in small vesicles (possibly endosomes) and then into vacuoles. As time progressed, more peptide accumulated in vacuoles causing these organelles to expand. They expanded to a point where they started occupying the fungal cells, and simultaneously, the shrinkage of nuclei was seen; this event coincided with cell death. It is interesting to note that a short synthetic hexapeptide, PAF26 (Munoz et al., 2012) and GMA4CG_V6 (Tetorya et al., 2023) were also localized into vacuoles and have the sequence motif WFW motif and cationic residues Arg and Lys in their sequences. Vacuoles are dynamic and multifunctional organelles responsible for ion homeostasis, detoxification and storage (Richards et al., 2010). Thus, one possibility is that the accumulation of GMAOe1C_V1* could trigger oxidative stress or alter the vacuole’s capability to maintain proper pH or ion homeostasis, leading to metabolic failure; however, the exact mechanism behind vacuolar expansion and cell death remains to be determined.

To gain further insight into the MoA of GMAOe1C_V1*, we investigated the global transcriptional response of *B. cinerea* to peptide challenge using RNA-seq, uncovering significant alterations in membrane integrity, oxidative stress regulation, amino acid biosynthesis, and core metabolic pathways. The cell wall serves as the primary site of interface for AMPs. The modulation of genes that are involved in reinforcement of the cell wall has also been shown to be a generalized cellular response to peptide-induced stress (López-García et al., 2010; Muñoz et al., 2013). A study by (Harries et al., 2013), specifically demonstrated the positive role of Eos1p, an enzyme involved in the N-linked glycosylation of proteins during PAF26 internalization in *S. cerevisiae.* The upregulation of several genes in our transcriptome analysis associated with the glycoside hydrolase (GH) family involved in degradation of cell wall polysaccharides and protein N-glycosylation suggests remodelling of cell wall upon peptide treatment which might be important for GMAOe1C_V1* internalization in germling cells. The pronounced overexpression of several membrane transporters suggests that the fungus may be striving to detoxify or expel the peptide through increased transmembrane efflux mechanisms across plasma membrane. This is consistent with previous studies that reported increased expression of ABC and MFS transporter genes in response to fungicide and peptide treatment in *B. cinerea* (Shi et al., 2020; Fan et al., 2024). Interestingly, the activation of genes encoding ubiquitin-related proteins, endopeptidases, proteases and unfolded protein was observed which could possibly be linked to the autophagy and subsequent cell death. However, none of the metacaspase or ATG genes were activated in response to treatment, suggesting that vacuolar expansion and cell death by GMAOe1C_V1* might be linked to metacaspase independent autophagic degradation. Similar observations by (Wang et al., 2023) reported that Perillaldehyde (PA), an antimicrobial natural compound inhibited *B. cinerea* growth by promoting ubiquitination and triggering autophagy-mediated protein degradation independent of metacaspases. This autophagy was related to disrupted mitochondrial function, ROS accumulation, alteration of intracellular Ca²⁺ levels in the pathogen upon PA treatment.

RNA-seq study identified an additional mechanism contributing to the antifungal activity of GMAOe1C_V1*, marked by the downregulation of essential mitochondrial function-related genes and amino acid biosynthesis pathways. Numerous AMPs have demonstrated the ability to disrupt mitochondrial gene expression, alter amino acid metabolism, and trigger the accumulation of ROS under peptide stress in plant pathogenic fungi including *B. cinerea* (Wang et al., 2023; Fan et al., 2024; Jin et al., 2024). Additionally, a number of antioxidant-related genes showed upregulation, which might be due to mitochondrial damage induced ROS in fungi, as previously documented for LpDef1 and Ppdef1 (Vieira et al., 2015; Parisi et al., 2024). Although transcriptome analysis provided insight into mitochondrial dysfunction, and oxidative stress, further work is necessary to clarify the structural abnormalities, particular mitochondrial disruptions, and their exact role in ROS accumulation. Overall, our findings indicate that GMAOe1C_V1* disrupts fungal homeostasis and leads to apoptosis by alterations of membrane related genes, enhancing oxidative stress responses, activating proteolytic pathways, and downregulating mitochondrial functions, thus exhibiting multimodal mechanisms of action. In support of *in vivo* translation inhibition via puromycin labeling, no transcripts encoding ribosomal proteins were detected in our transcriptome data. Thus, it is likely GMAOe1C_V1* affects cellular homeostasis through pathways associated directly or indirectly with synthesis of proteins rather than direct interaction with ribosomes or interfering with the cell cycle. In summary, our study indicated multi-faceted antifungal action of GMAOe1C_V1* peptide and demonstrated its potential for development as a biofungicide.

## MATERIALS AND METHODS

### Fungal cultures, growth medium, and spore collection

Glycerol stock maintained at -80°C was used for the revival of fungal strain *B. cinerea* T4 on V8 medium at 25°C. Three-week-old *B. cinerea* spores were harvested by flooding the fungal plates with sterile water. The spore suspension was filtered through two layers of Miracloth, centrifuged at 13000 rpm for 2 min, and washed 2-3 times using sterile water. Fungal conidia were resuspended in low-salt synthetic fungal medium (SFM) (Liang et al., 2001). The pathogen suspension was adjusted to the desired spore density using a hemocytometer.

### Purification of OefDef1.1 and chemical synthesis of OefDef1.1-derived peptides

The *Pichia pastoris* X33 strain expressing the full-length OefDef1.1 was used for purification of this defensin as previously described (Li et al., 2019). The 22-amino acid OefDef1.1(G32-Y53), designated GMAOe1C, and its variants GMAOe1C_V1 and GMAOe1C_V2 with and without a C3-C21 disulfide bond were designed by chemical synthesis. Chemically synthesized peptides with greater than 90% purity were obtained from WatsonBio Sciences. All peptides were resuspended in nuclease-free water and their concentrations were determined using the bicinchoninic acid (BCA) assay.

### *In vitro* antifungal activity assays & alanine scanning mutagenesis

Antifungal activity of peptides against *B. cinerea* was determined using a 96-well plate resazurin cell viability assay (Chadha and Kale, 2015; Li et al., 2019). A change in the color of resazurin from blue to pink or colorless indicated the presence of live fungal cells. The concentration at which no change in color was observed was considered as minimal inhibitory concentration (MIC) value for each peptide. Fungal growth inhibition was also observed under the microscope and images were captured using Leica CMI6000B microscope with 20X objective lens. Antifungal activity of peptides was also tested in SFM supplemented with cations. Four chemically synthesized alanine scanning variants of GMAOe1C_V1, each without a disulfide bond, were tested for *in vitro* antifungal activity against *B. cinerea* and MIC of each variant was determined using the resazurin cell viability assay.

### Circular dichroism

For circular dichroism (CD) studies, OefDef1.1, GMAOe1C_WT and its variants were resuspended in MilliQ water at a concentration of 0.2 mg/mL. All measurements were performed on an Applied Photophysics Chirascan instrument at 20°C using a 2 mm quartz cell. The spectra presented are the average of three independent scans at 1 nm resolution (0.5 seconds per wavelength) between 200 and 300 nm. Data were collected using the Chirascan software (version 4.2.12) and visualized using custom Python scripts. Molar ellipticity [θ] (y-values) was calculated from the raw CD values (‘A’) using sample concentration (0.2 mg/mL) (‘c’), the peptide molecular weight (variable) (‘M’), and the path length (0.2 cm) (‘L’) with the following relation:

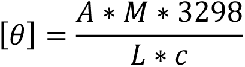

The BeStSel web tool (https://bestsel.elte.hu/index.php) was used to fit the experimental data and estimate relative contributions from different secondary structure elements (Micsonai et al., 2015; Micsonai et al., 2018; Micsonai et al., 2022).

### Structure Prediction

Structures were predicted using the AlphaFold2 (AF2) colab implementation (https://colab.research.google.com/github/sokrypton/ColabFold/blob/main/AlphaFold2.ipynb). The highest-ranked structure based on pLDDT scores was used as a representative structure. All models were visualized and all graphics were generated using ChimeraX (Goddard et al., 2018). The AF2 PDB model of OefDef1.1 was used to calculate a theoretical CD spectrum using the computational method, SESCA (Nagy et al., 2019).

### SYTOX Green membrane permeabilization assay

Time-lapse laser scanning confocal microscopy was performed to monitor *B. cinerea* membrane permeabilization by GMAOe1C_V1using the membrane impermeant SYTOX Green (SG) dye. Fresh conidia were isolated and geminated in glass-bottom petri dishes (MatTek Corporation, Ashland, MA) for 10-12 h in 1X SFM medium. Subsequently, germlings were mixed with GMAOe1C_V1 at sub-lethal dose-1.5 μM and 1 μM SG dye and analyzed for membrane permeabilization using Leica SP8-X confocal microscopy. Fluorescence was observed at excitation and emission wavelengths of 504 nm and 515-590 nm, respectively.

### Internalization and subcellular localization of GMAOe1C_V1* in *B. cinerea*

Time-lapse confocal laser scanning microscopy was performed to monitor the internalization and subcellular localization of tetramethyl rhodamine (TAMRA)-labeled GMAOe1C_V1* in *B. cinerea* germlings. For subcellular localization studies, a sub-lethal (3 μM) and lethal dose (6 μM) of TAMRA-labeled peptide was added to *B. cinerea* germlings along with the vacuole-specific dye, 7-amino-4-chloromethylcoumarin (CMAC, 10 μM) or nucleus-staining dye, Hoechst33258 (100 μg/mL). The excitation and emission wavelengths of TAMRA labeled-GMAOe1C_V1* are 552 nm and 565-600 nm, for CMAC and Hoechst33258 405 nm and 416-484 nm, respectively. In addition, propidium iodide (PI, 15 µM), a cell death marker was used to determine the cell death timing of GMAOe1C_V1 (3 µM) challenged *B. cinerea* germlings at an excitation and emission wavelength of 493 nm and 560-646 nm, respectively. A z-stack was acquired at each time point using a 63X water immersion objective. All images were processed using LASX software.

### *In vitro* translation inhibition assay

Total RNA was extracted and purified from *B. cinerea* germlings grown for 20 h in Potato Dextrose Broth (Difco^TM^, Fisher Scientific, USA) using the NucleoSpin RNA Plus Mini kit (Takara Bio, USA) according to the manufacturer’s instructions. Gel shift experiments were performed by mixing 200 ng of the total fungal RNA with different concentrations of GMAOe1C_V1* in 20 μL of binding buffer (10 mM Tris-HCl, pH 8.0, 1 mM EDTA). Reaction mixtures were incubated for 1 h at room temperature and run on a 1.2% agarose gel. RNA in the agarose gel was visualized using a Bio-Rad ChemiDoc XRS + system. *In vitro* translation was performed using the RiboMAX Large Scale RNA Production System and the Wheat Germ Extract *in vitro* translation system (Promega, WIS). The luciferase DNA template was translated to luciferase following the manufacturer’s protocol, and the luciferase activity was assessed using luminometer (Tecan). Different concentrations of GMAOe1C_V1* were used to determine the translation inhibitory activity.

### *In vivo* translation inhibition assay using puromycin labeling

Puromycin labeling was used to evaluate protein translation inhibition by GMAOe1C_V1* in *B. cinerea* germling cells (Godwin and Shah, 2025). *B. cinerea* spores were grown at a concentration of 1 × 10^5^ conidia/mL in 1x SFM for 24 h at 28°C with shaking at 220 rpm. The resulting germlings were harvested by centrifugation at 4000 rpm for 15 min and then resuspended in a 1X SFM solution (2 mL) containing GMAOe1C_V1* at a concentration of 1.5 or 3 µM, water (negative control), or cycloheximide (positive control) at 100 μg/mL concentration. Germlings were incubated for 1 h at 28°C with shaking at 220 rpm and gently washed thrice with water and harvested at 4000 rpm for 15 min and resuspended in fresh 1x SFM (2 mL). To assess protein translation, puromycin (1 mM) was added, and the cells were incubated for 2 h at 28°C with shaking. After incubation, the cells were washed 2-3 times with water, and frozen in liquid N_2_. Proteins were extracted using the isolation buffer and concentrations were determined using the BSA assay. Next, equal amounts of protein were separated on a 4-16% SDS-PAGE gradient gel, followed by transfer onto a nitrocellulose membrane. The membrane was placed into the blocking solution (3% bovine serum albumin prepared in PBST) for 1 h followed by incubation in primary antibody against puromycin (MABE343, Sigma-Aldrich) at a 1:3000 dilution of the blocking solution to detect puromycin incorporation. The blot was washed and incubated with secondary antibody (1:15000) for 1 h and probed using SuperSignal™ West Pico PLUS Chemiluminescent Substrate (Godwin and Shah, 2025; Pokhrel et al., 2025).

### Antifungal assays using detached pepper leaves

Antifungal activity of peptides was compared on detached leaves from 4-week-old pepper (*Capsicum annuum*, California Wonder 300 TMR) plants as described previously (Tetorya et al., 2023). Each peptide was tested at a concentration of 1.5, 3 and 6 μM. *B. cinerea* spores (5 x 10^5^/mL in SFM) and peptide in equal volumes were applied at the same spots and incubated in a humid chamber in dark. The lesions were photographed 2 days post-inoculation in white light and CropReporter (Phenovation). The relative lesion sizes were determined using ImageJ software.

### *In planta* post-inoculation antifungal activity assays

The post-inoculation antifungal activity of peptides against *B. cinerea* was tested on four-week-old pepper plants grown under controlled growth conditions (14 h light/10 h dark cycles). Each peptide was tested at concentrations of 6 and 12 μM against *B. cinerea*. Pepper plants were spray-inoculated with 1 mL of 5 x 10^5^ spores/mL. After 2 h, GMAOe1C_WT, GMAOe1C_V1* and GMAOe1C_V2* peptides were sprayed onto inoculated leaves. Plants were observed for gray mold disease symptoms at 3 dpi and photographed. Disease severity index was also calculated based on visual symptoms and scored (Segarra et al., 2007).

### RNA-seq analysis

Ten mL of *B. cinerea* conidia at a concentration of 1 × 10^5^/mL in 1x SFM were incubated for 24 h at 28°C with shaking at 220 rpm. Peptides were added and germlings were collected at 30 min and 1 h. Germlings collected at 0 h were used as a control with no peptide treatment. The samples were centrifuged, immediately frozen in liquid nitrogen and stored at -80°C. Total RNA was extracted from pretreated *B. cinerea* germlings (control) and GMAOe1C_V1* treated germlings at 30 min and 1 h using Quick-RNA^TM^ MiniPrep Plus kit following manufacturer’s guidelines. The integrity of RNA was checked by 1.2% agarose gel and Nanodrop 2000c (Spectrophotometer, Thermo Scientific). After quality check, RNA sequencing was performed by Novogene Corporation using Illumina NovaSeq X Plus for a total of nine samples consisting of three biological replicates (A, B, C) from each treatment (0 h, 30 min, 1 h). The clean raw sequencing reads were mapped and aligned to the reference genome of *B. cinerea* B05.10 (ensembl_42_botrytis_cinerea_asm83294v1). Nine libraries were constructed with each treatment having three biological replicates denoted as A, B and C. The detailed summary statistics are provided in Supplementary **Table S1**. The gene counts were further taken as an input and differential expression analysis was conducted using DESeq2 v1.24 package. Differentially Expressed Genes (DEGs) with a threshold FDR <= 0.05 and log_2_FC >1 or <-1 were generated from the pairwise comparison of the treated *B. cinerea* germlings at both time points (30 min and 1 h) with 0 h as the control sample. Variance stabilizing transformation (VST) was applied to normalize the count data in DESeq2 and was utilized for exploratory data analysis PCA plot, hierarchical clustering and heatmaps.

The statistically significant DEGs were used for functional enrichment analysis. Functional classification, GO enrichment and pathway enrichment analysis was performed using the DAVID tool (https://davidbioinformatics.nih.gov) (Sherman et al., 2022) and ShinyGO version v0.82 (http://bioinformatics.sdstate.edu/go/) (Ge et al., 2020). Pathway enrichment was conducted using the KEGG database (https://www.genome.jp/kegg/). The enrichment dot plots were constructed in R with ggplot2 package.

### RT-qPCR for validation of RNA-seq

Total RNA was isolated from Quick-RNA MiniPrep Plus as per manufacturer’s instructions. 500 ng of RNA was used for first-strand cDNA synthesis using the RevertAid First Strand cDNA Synthesis kit (ThermoFisher Scientific). The product of the first-strand cDNA synthesis as template and primer pairs designed using Primer3Plus (Supplementary **Table S2)** were used to perform RT-qPCR following the instructions provided by the igSYBR Green qPCR 2X Master Mix kit. For relative gene expression analysis, the 2^−ΔΔCt^ method (Livak and Schmittgen, 2001) was used to calculate the expression of genes relative to that of the GAPDH gene (*Bcin15g02120*).

### Statistical analysis

All experiments were performed in two or more biological replicates. Plots were generated and statistical tests were run using R4.2.2. The comparison of p-value ≤ 0.05 was considered statistically significant.

## Acknowledgements

We thank Dr. Hui Li for providing the *Pichia pastoris* isolate expressing OefDef1.1. We are grateful to Dr. Ambika Pokhrel for providing her critical comments on the manuscript and suggestions for improving it. We are grateful to Dr. Kirk Czymmek for providing his guidance during the confocal microscopy experiments. We acknowledge Advanced Bioimaging Laboratory 690 (RRID: SCR_018951) at the Donald Danforth Plant Science Center (DDPSC) for imaging support and the use of the Leica SP8-X confocal microscope funded by an NSF Major Research Instrumentation grant (DBI-1337680). The use of phenotyping, greenhouse, and growth chamber facilities provided by the DDPSC is acknowledged.

## Author contributions

D.S. conceived the idea, supervised, reviewed and edited the manuscript. R.T., H.S., J. G., M.T. and E.U. designed, performed the experiments, analyzed the data, and wrote the manuscript. D.S., M.T. designed the peptide sequences. All authors have reviewed and edited the manuscript.

## Data availability

The sequence data supporting this study have been deposited in NCBI’s SRA database under the accession number PRJNA1356523.

## Supplementary Figures

**Fig. S1.** Circular dichroism analysis and structure prediction by AlphaFold2 Measured and predicted structural features for OefDef1.1. **A)** Measured CD spectrum (solid line) of OefDef1.1 compared to that calculated from an AlphaFold2 (AF2) model (dashed line) of the protein. **B)** AF2 rank 1 model colored according to per-residue pLDDT score (blue = pLDDT > 90; cyan = pLDDT > 70). **C)** Surface representation of the rank 1 AF2 model colored to reflect electrostatic potential (red = negative; blue = positive). The sequence of OefDef1.1 (MW = 6207.17 Da) is shown in Fig. 1A.

**Fig. S2.** Structural features of OefDef1.1-derived peptides. **A)** CD spectrum of the peptide GMAOe1C_WT and AF2-predicted structure which shows poor confidence by pLDDT scores. **B)** CD spectrum of the GMAOe1C_V1* and its AF2-predicted structure. **C)** CD spectrum of the GMAOe1C_V2* and its AF2-predicted structure. All AF2 models are colored according to pLDDT score (blue = pLDDT > 90; cyan = pLDDT > 70; yellow = pLDDT > 50; orange = pLDDT < 50). The sequences of GMAOe1C_WT; GMAOe1C_V1* and GMAOe1C_V2* are shown in Fig. 1A.

**Fig. S3.** Antifungal activity of OefDef1.1 and OefDef1.1-derived peptides against *B. cinerea* on detached pepper leaves. **A)** Schematic representation of the peptides and spores alone applied on the surface of pepper leaves. **B-D)** Different concentrations of peptides were applied, followed by spore addition at a concentration of 5 x 10^4^ spores/mL at each spot. Gray mold disease lesions at 3 dpi were imaged under white light and high-resolution images were taken using CropReporter. **E-G)** Lesion sizes were measured using ImageJ, and the relative lesion sizes are shown as mean±SD. One-way Anova with Tukey’s multiple comparison test was applied. ns-p>0.05, *p<0.05, **p<0.01, ***p<0.001, *p<0.0001.

**Fig. S4.** Confocal images showing GMAOe1C_V1*-induced cell death in *B. cinerea* germlings using propidium iodide (PI) staining. GMAOe1C_V1 (3 μM) was used to determine the cell death timings of *B. cinerea* germlings. Bright field shows that the vacuole expands (T = 20 min) and the germling tip shrinks (T = 20 min). With time shrinkage of cells increases towards the germling conidial head (T = 30 min).

**Fig S5.** Gel retardation assay to reveal the interaction of GMAOe1C_V1*with *B. cinerea* total RNA. Gel retardation assay revealed that GMAOe1C_V1* does not bind to ribosomal RNA at peptide concentrations used. Total RNA was incubated with different concentrations of the GMAOe1C_V1* peptide, and RNA was visualized on a 1.2% agarose gel. Sterile water was used as a negative control and NCR13_PFV1 peptide (Godwin et al., 2024) was used as positive control.

**Fig. S6.** Transcriptomic reprogramming upon GMAOe1C_V1* peptide treatment. **A)** Diagrammatic representation of experimental design for RNA-Seq analysis **B)** Principal Component analysis (PCA) showing distinct gene expression profiles of control (0 h) and treated (30 min and 1h) samples along PC1 (40%). **C)** Venn diagrams showing number of exclusive and shared DEGs upregulated or downregulated following peptide treatment.

**Fig. S7.** MA scatter plot depicting DEGs (blue dots) based on their expression levels across samples (x-axis) and log2fold change (y-axis). The plots show all the DEGs before **(A, C)** and after **(B, D)** filtration of p-adjusted value at 30 min and 1 h post-treatment with respect to the control samples. The blue dots represent upregulated and downregulated DEGs while the grey dots represent genes with no significant difference.

**Fig. S8.** Validation of RNA-seq-identified DEGs using qPCR. **A-H)** Expression levels of upregulated and downregulated genes. GAPDH has been used as an internal control. The statistical significance analysis was performed using one-way ANOVA with Tukey multiple comparisons test. Error bars denote Mean±SEM.

**Fig. S9.** KEGG enrichment **(A, B)** and Gene ontology (GO) enrichment analysis **(C)** of all downregulated DEGs (132) in response to GMAOe1C_V1* treatment.

**Fig. S10.** Summary of various metabolic pathways (KEGG: bfu01100) which might be involved in antifungal mechanism in *B. cinerea* in response to GMAOe1C_V1* peptide treatment

**Fig. S11.** Schematic representation of various genes involved in biosynthesis of secondary metabolites misregulated upon GMAOe1C_V1* peptide treatment at both time points. The downregulated genes are shown in red color, and upregulated genes are shown in blue color.

## Supplementary Tables

**Table S1**. Summary statistics of transcriptome data for all nine samples

**Table S2**. Primer list for quantitative PCR

**Table S3**. Description of genes used for validation of RNA-seq-identified DEGs using qPCR

**Table S4**. Gene ontology and KEGG enrichment of DEGs at each time point (30 min and 1 h) post GMAOe1C_V1* peptide treatment with respect to 0 h (control)

**Table S5**. Expression of upregulated DEGs involved in various processes and pathways

**Table S6**. Expression of downregulated DEGs involved in various processes and pathways

## Supplementary videos

**Video 1**. Uptake of SYTOX dye in the presence of GMAOe1C_V1*

**Video 2**. Colocalization of FM4-64 and TAMRA-GMAOe1C_V1*

**Video 3**. Internalization of TAMRA-GMAOe1C_V1*

**Video 4**. Colocalization of Hoechst and TAMRA-GMAOe1C_V1*

**Video 5**. Colocalization of CMAC and TAMRA-GMAOe1C_V1*

Supplementary videos can be accessed at https://tinyurl.com/bdh8cjrc

