## Supplementary figures for "Modes of action and *in planta* antifungal activity of *Olea europaea* defensin OefDef1.1-derived peptide variant"

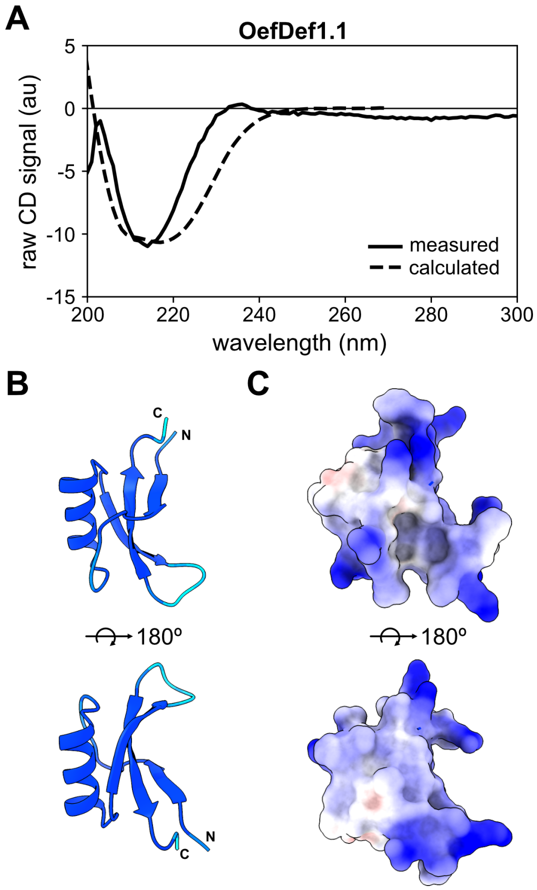


**Fig S1. Circular dichroism analysis and structure prediction by AlphaFold2** Measured and predicted structural features for OefDef1.1. **A)** Measured CD spectrum (solid line) of OefDef1.1 compared to that calculated from an AlphaFold2 (AF2) model (dashed line) of the protein. **B)** AF2 rank 1 model colored according to per-residue pLDDT score (blue = pLDDT > 90; cyan = pLDDT > 70). **C)** Surface representation of the rank 1 AF2 model colored to reflect electrostatic potential (red = negative; blue = positive). The sequence of OefDef1.1 (MW = 6207.17 Da) is shown in Fig. 1A.

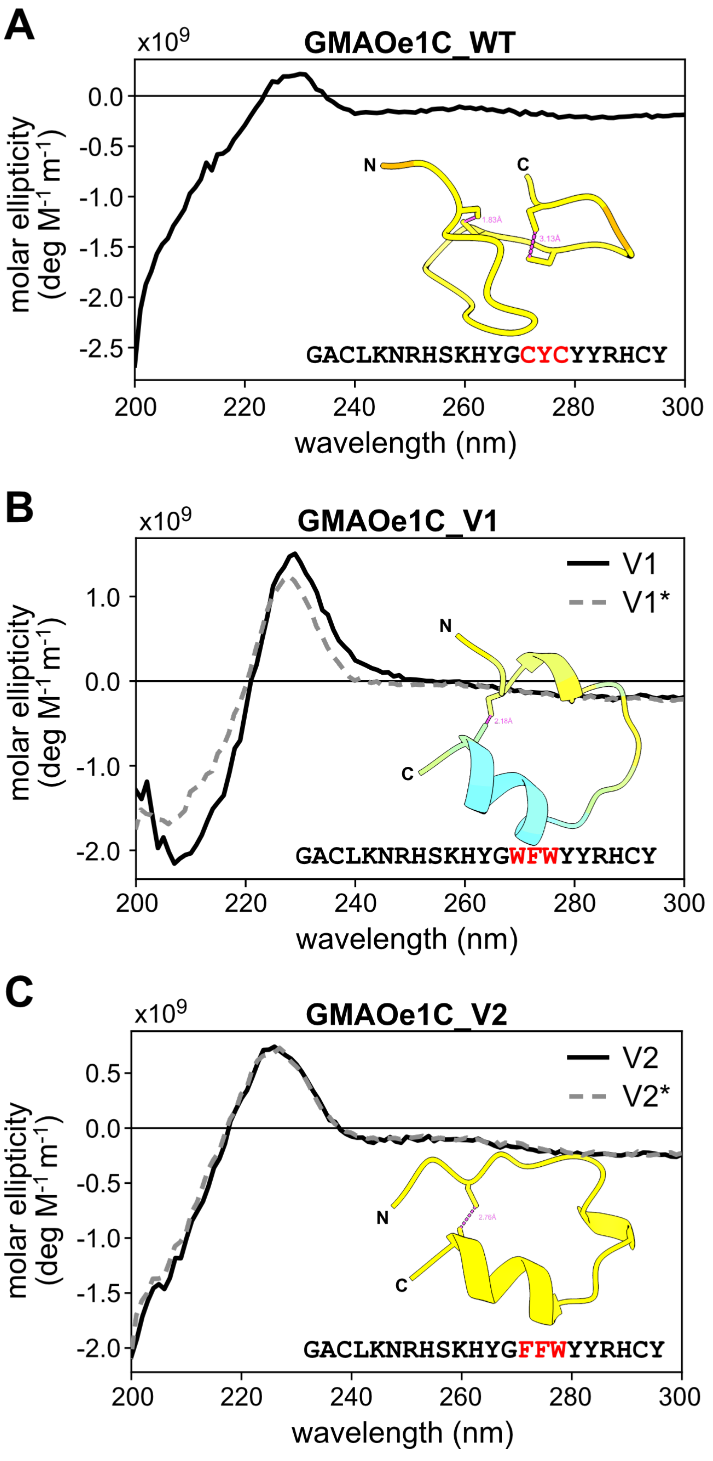


**Fig S2.** Structural features of OefDef1.1-derived peptides. **A)** CD spectrum of the peptide GMAOe1C_WT and AF2-predicted structure which shows poor confidence by pLDDT scores. **B)** CD spectrum of the GMAOe1C_V1* and its AF2-predicted structure. **C)** CD spectrum of the

GMAOe1C_V2* and its AF2-predicted structure. All AF2 models are colored according to pLDDT score (blue = pLDDT > 90; cyan = pLDDT > 70; yellow = pLDDT > 50; orange = pLDDT < 50). The sequences of GMAOe1C_WT; GMAOe1C_V1* and GMAOe1C_V2* are shown in Fig. 1A.

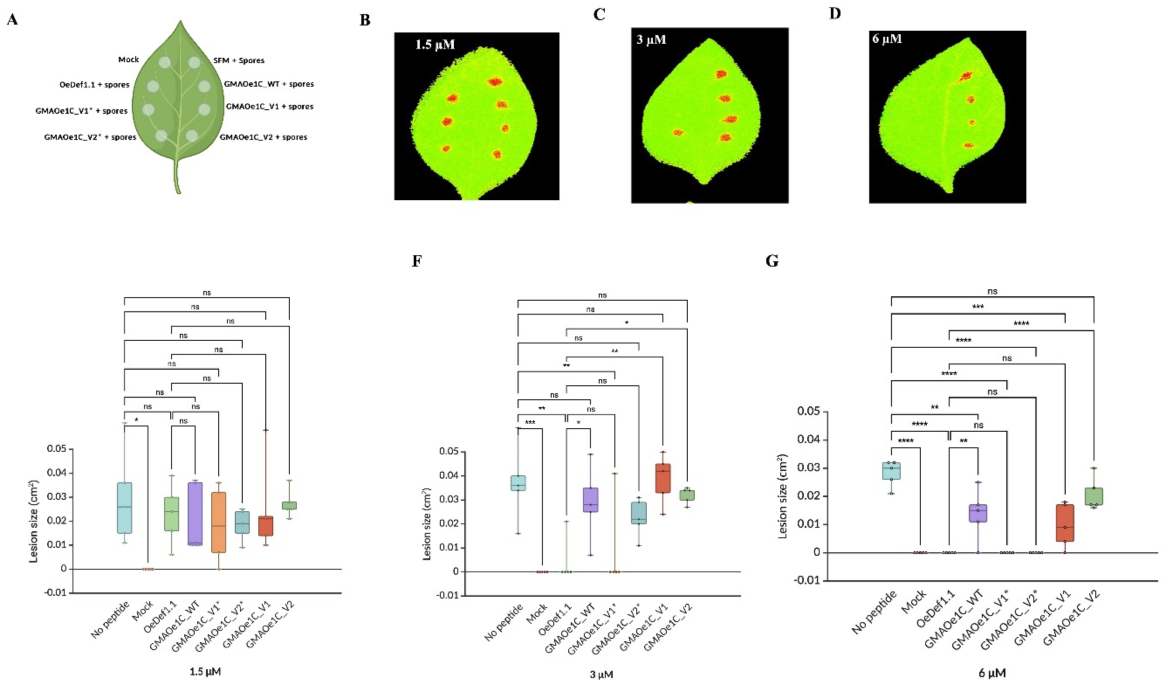


**Fig S3.** Antifungal activity of OefDef1.1 and OefDef1.1-derived peptides against *B. cinerea* on detached pepper leaves. **A)** Schematic representation of the peptides and spores alone applied on the surface of pepper leaves. **B-D)** Different concentrations of peptides were applied, followed by spore addition at a concentration of 5 x $10^4$ spores/mL at each spot. Gray mold disease lesions at 3 dpi were imaged under white light and high-resolution images were taken using CropReporter. **E-G)** Lesion sizes were measured using ImageJ, and the relative lesion sizes are shown as mean±SD. One-way Anova with Tukey's multiple comparison test was applied. ns- p>0.05, *p<0.05, **p<0.01, ***p<0.001, *p<0.0001.

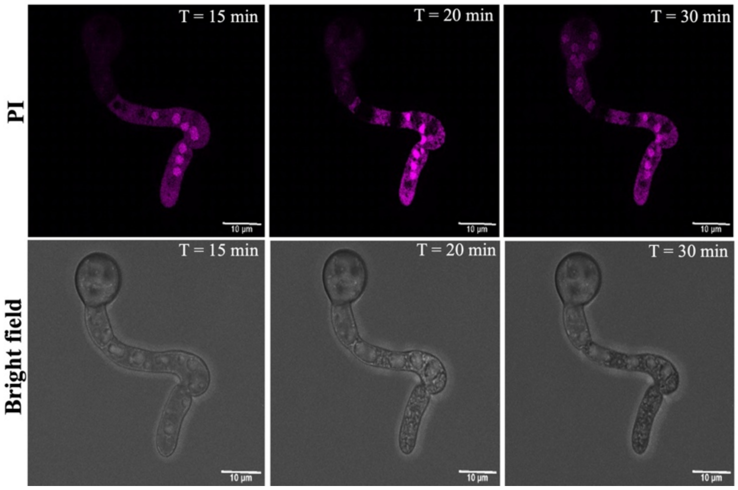


**Fig S4.** Confocal images showing GMAOe1C_V1*-induced cell death in *B. cinerea* germlings using propidium iodide (PI) staining. GMAOe1C_V1 (3 µM) was used to determine the cell death timings of *B. cinerea* germlings. Bright field shows that the vacuole expands (T = 20 min) and the germling tip shrinks (T = 20 min). With time shrinkage of cells increases towards the germling conidial head (T = 30 min).

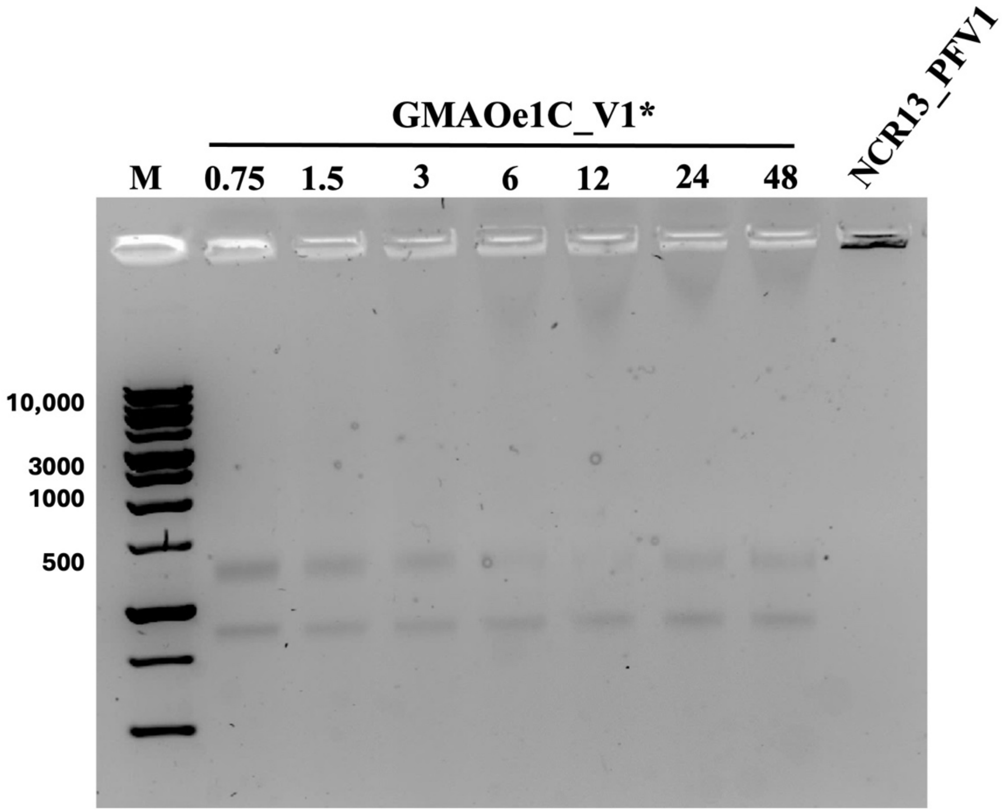


**Fig S5.** Gel retardation assay to reveal the interaction of GMAOe1C_V1*with *B. cinerea* total RNA. Gel retardation assay revealed that GMAOe1C_V1* does not bind to ribosomal RNA at peptide concentrations used. Total RNA was incubated with different concentrations of the GMAOe1C_V1* peptide, and RNA was visualized on a 1.2% agarose gel. Sterile water was used as a negative control and NCR13_PFV1 peptide (Godwin et al., 2024) was used as positive control.

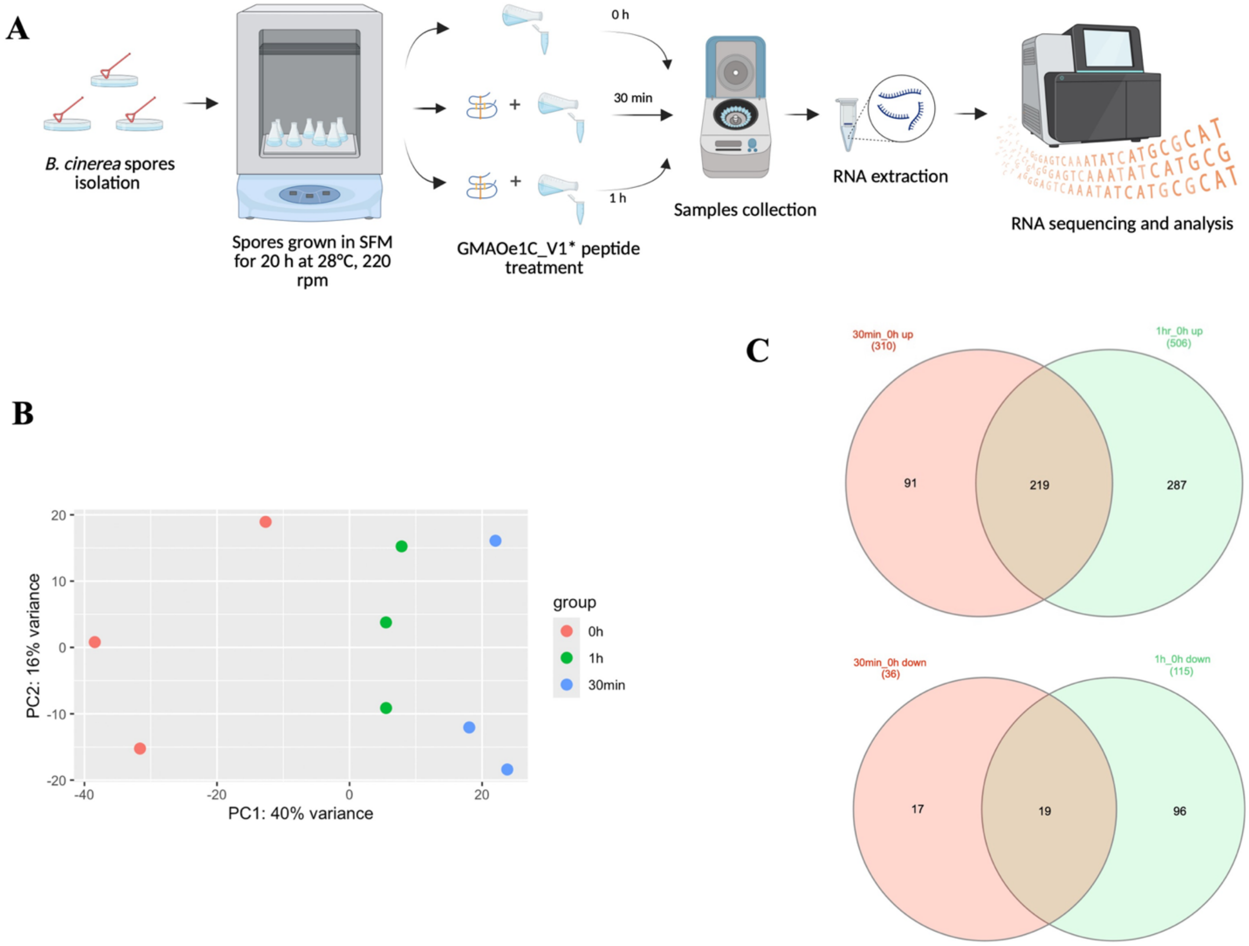


**Fig S6.** Transcriptomic reprogramming upon GMAOe1C_V1* peptide treatment. **A)** Diagrammatic representation of experimental design for RNA-Seq analysis **B)** Principal Component analysis (PCA) showing distinct gene expression profiles of control (0 h) and treated (30 min and 1h) samples along PC1 (40%). **C)** Venn diagrams showing number of exclusive and shared DEGs upregulated or downregulated following peptide treatment.

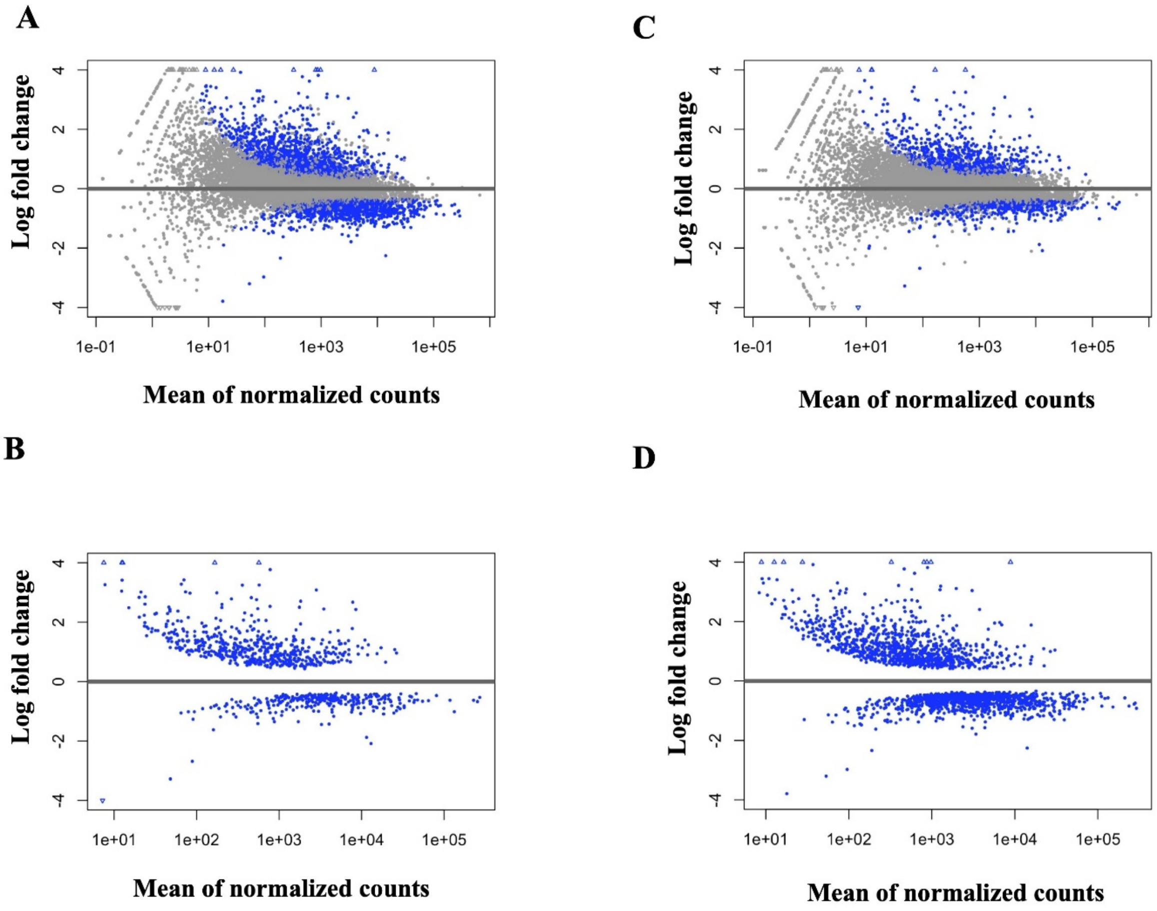


**Fig S7.** MA scatter plot depicting DEGs (blue dots) based on their expression levels across samples (x-axis) and log2fold change (y-axis). The plots show all the DEGs before **(A, C)** and after **(B, D)** filtration of p-adjusted value at 30 min and 1 h post-treatment with respect to the control samples. The blue dots represent upregulated and downregulated DEGs while the grey dots represent genes with no significant difference.

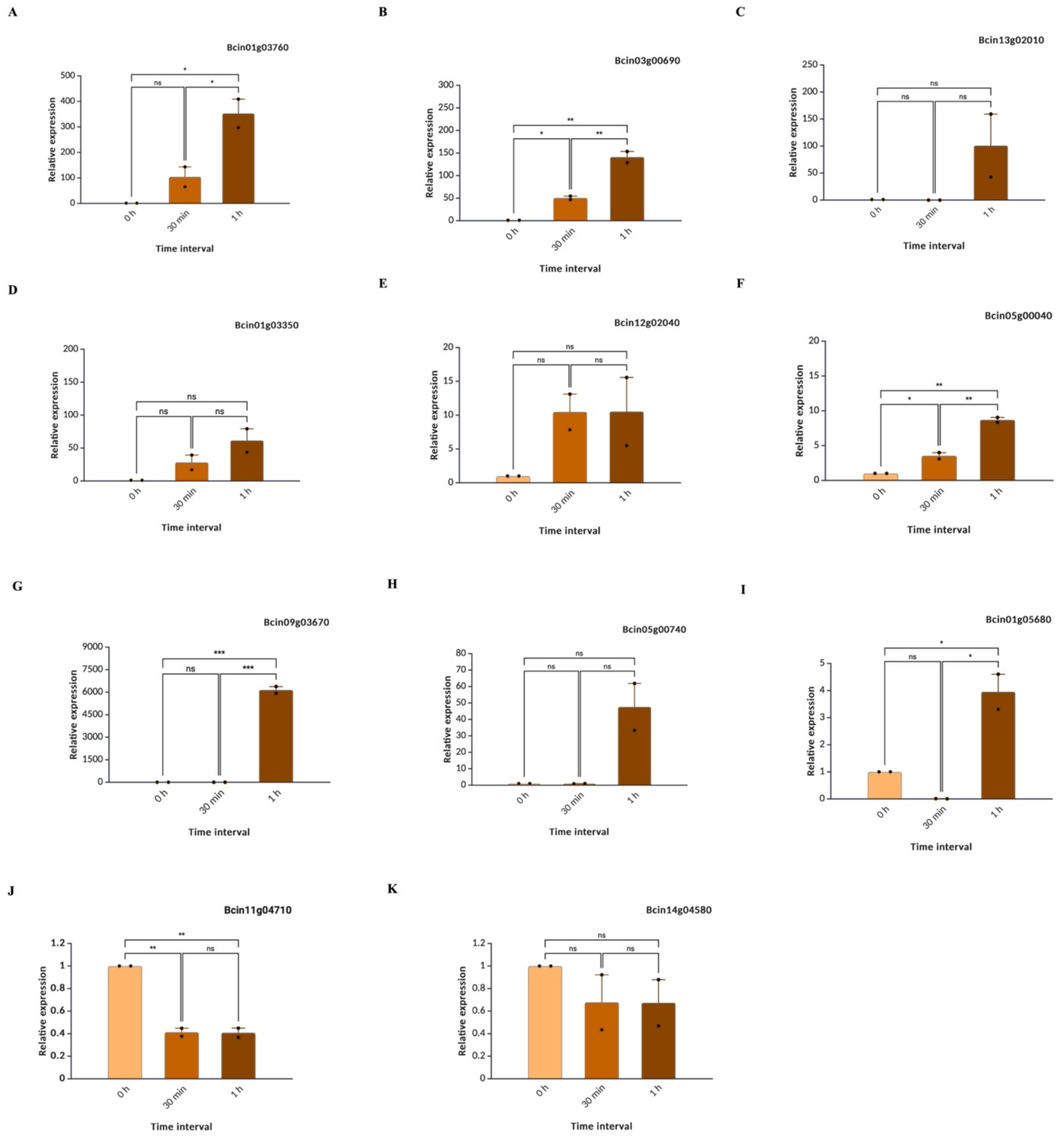


**Fig S8.** Validation of RNA-seq-identified DEGs using qPCR. **A-H)** Expression levels of upregulated and downregulated genes. GAPDH has been used as an internal control. The statistical significance analysis was performed using one-way ANOVA with Tukey multiple comparisons test. Error bars denote Mean±SEM.

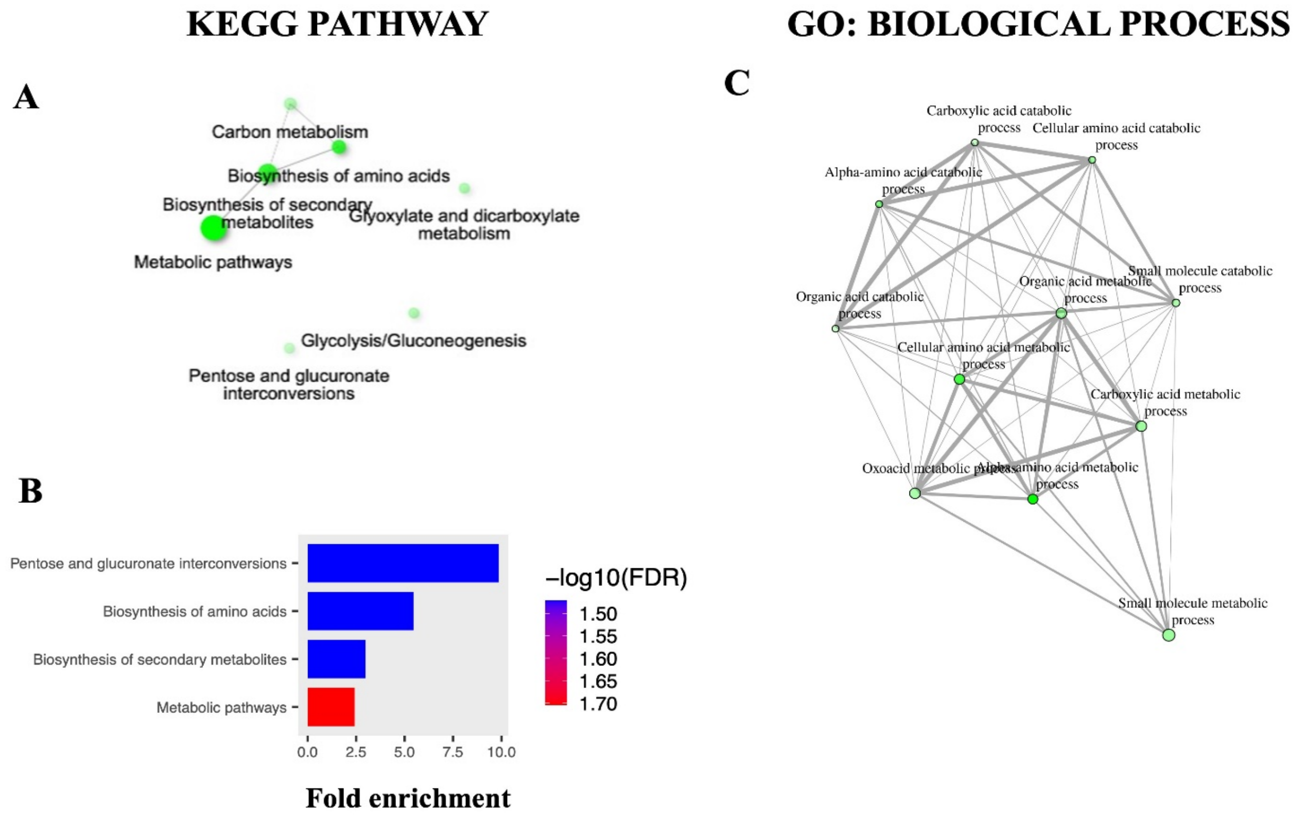


**Fig S9.** KEGG enrichment **(A, B)** and Gene ontology (GO) enrichment analysis **(C)** of all downregulated DEGs (132) in response to GMAOe1C_V1* treatment.

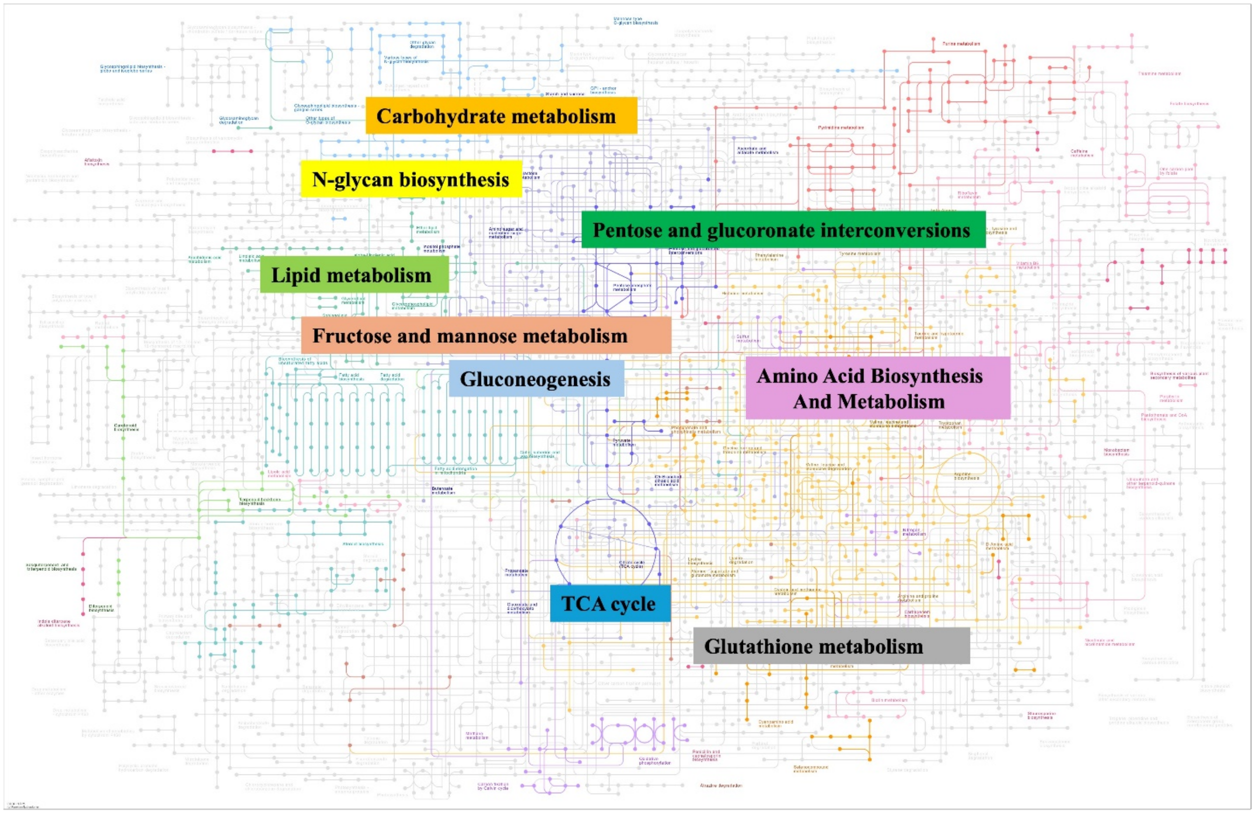


**Fig S10.** Summary of various metabolic pathways (KEGG: bfu01100) which might be involved in antifungal mechanism in *B. cinerea* in response to GMAOe1C_V1* peptide treatment.

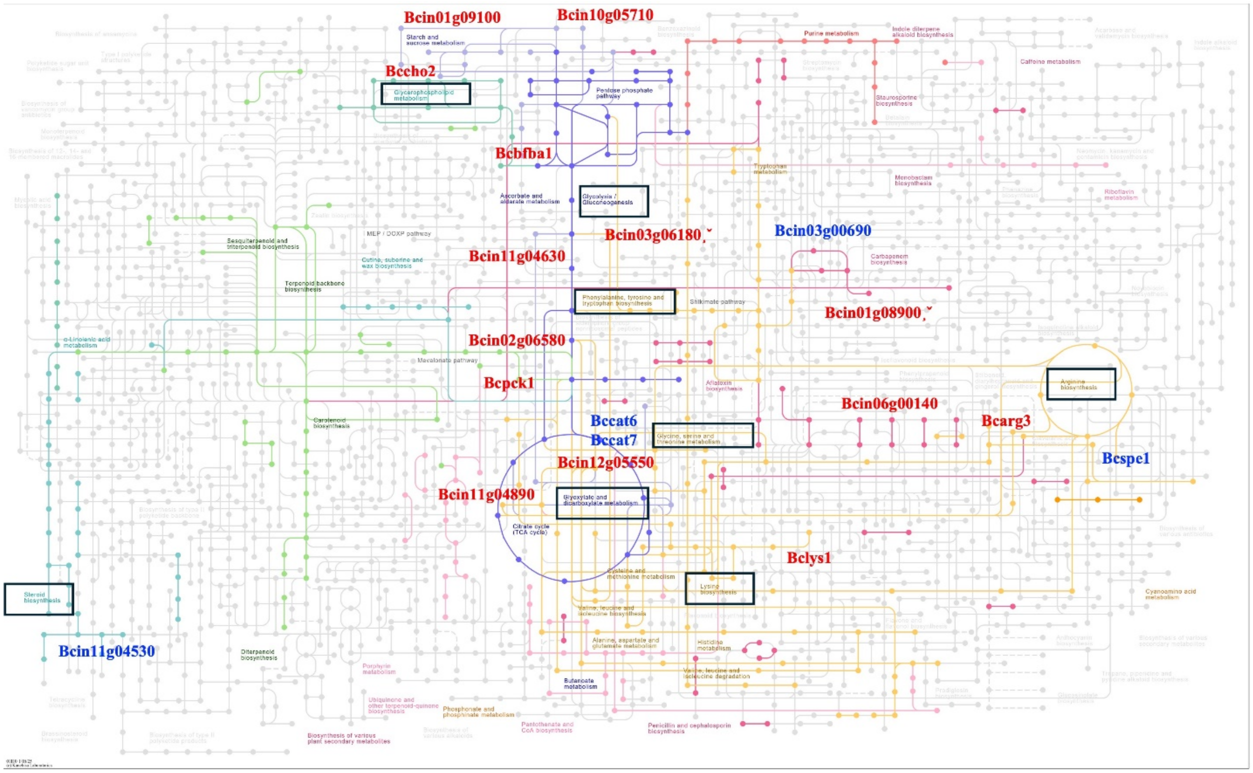


**Fig S11.** Schematic representation of various genes involved in biosynthesis of secondary metabolites mis regulated upon GMAOe1C_V1* peptide treatment at both time points. The downregulated genes are shown in red color, and upregulated genes are shown in blue color.
