## Supplementary Table S2 for "Modes of action and *in planta* antifungal activity of *Olea europaea* defensin OefDef1.1-derived peptide variant"

| **Bcin12g05930.1** | Forward primer: CTCAAGGCACTCTACTCGCC  Reverse primer: GCCAGGAGGACGGTATTTCC |
| --- | --- |
| **Bcin12g05920.1** | Forward primer: GAGGTGTGATGGGAGACAGC  Reverse primer: CTGGTTCTGGATCGCGACTT |
| **Bcin12g05950.1** | Forward primer: AACACGCTCTATCATCCCCG  Reverse primer: TTTCGCAGGAGTTGATGGCT |
| **Bcin12g05940.1** | Forward primer: CCGGCGAAAACAGGTCTACT  Reverse primer: GACCTTTAGCCTCATGCCGT |
| **Bcin14g04580.1** | Forward primer: CGGAACCGATTTGCACATCC  Reverse primer: CAGCTACAACTCTGTCGCCA |
| **Bcin11g04710.1** | Forward primer: ACACAGCTCTTGCATACGGA  Reverse primer: TGCCATGGGTTGTCGAGTTT |
| **Bcin09g00930.1** | Forward primer: CTCCGAATGTCTCCAACCCC  Reverse primer: CGTGGTAGATGTCTCGACCG |
| **Bcin03g05820.1** | Forward primer: CTCCAACCCCAACCTCTGAC  Reverse primer: CACCGTTGGTACTGGCGTAA |
| **Bcin13g05420.1** | Forward primer: TCCGGAATAAAGAATTCGCCA  Reverse primer: AACTCCTGCTGCAGTAAGTG |
| **Bcin08g00430.1** | Forward primer: GCGGGTGATGTATGGCGTTA  Reverse primer: TGCTGATGTAGAAAGGTTCAAGC |
| **Bcin13g01890.1** | Forward primer: CGCTTTCCCAGACAGCTACA  Reverse primer: GATGAACCCTGTGCCCAAGA |
| **Bcin12g02040.1** | Forward primer: GTTGACAAAGGACATGCCCG  Reverse primer: GGGCAACGTAGGAGACATCC |
| **Bcin09g02930.1** | Forward primer: GAGCCCACTAAACCTGGACC  Reverse primer: CTGGGACTTCACCGTTCTCC |
| **Bcin02g03130.1** | Forward primer: GACTCCCGAATCCCTGCAAA  Reverse primer: CCAACAGCTTTAACAGTGCGA |
| **Bcin13g00600.1** | Forward primer: TCGTCGCCTCCTTAGCTCTA  Reverse primer: TTGGTGCTGGTAGTCGAACC |
| **Bcin13g02010.1** | Forward primer: TTCGCTCTTGTCTCCTCTGC  Reverse primer- ACGGTATCACCAACTTGAACAGT |
| **Bcin09g03670.1** | Forward primer: AGTCGAAGCTGTGGAACTCG  Reverse primer: TCAGAACGGGCGTTTTTCCT |
| **Bcin01g05680.1** | Forward primer: TAACTACGTTGGCTGCACCC  Reverse primer: GAGGGAAGGCGTGTTGGTAA |
| **Bcin05g00040.1** | Forward primer: CCTCGAAGTCGCACACAGTT  Reverse primer: AATGATGCATTCTGCTCCGC |
| **Bcin01g03760.1** | Forward primer: CATCCCAACCATCCTCGCTC  Reverse primer: GAGCCGCGGATGTTGTAGTA |
| **Bcin03g00690.1** | Forward primer: TACGAAAAAGCCCTCCGTGA  Reverse primer: TCGAAAACGCTGGATTTCCG |
| **Bcin05g00740.1** | Forward primer: CAGTTAGCCTTGCAACAGCG  Reverse primer: AACATAGGAGGCAACGGAGC |
| **Bcin01g03350.1** | Forward primer: CACTATACACCTCCCGTCGC  Reverse primer: TGAAGTGTGTTGAGAGGCCG |
| **UBQ** | Forward primer: CAAGGTTACCGACAACAATA  Reverse primer: GCATCCATCAACTTCTTCAA |
| **UCE** | Forward primer: ATCACCCAAACATCAACT  Reverse primer: CATAGAGCAGATGGACAA |
| **GAPDH** | Forward primer: TGCCAAGAAGGTTGTTATC  Reverse primer: TGTAGGTCTCGTTGTTGA |
